# Gene regulatory network inference from CRISPR perturbations in primary CD4+ T cells elucidates the genomic basis of immune disease

**DOI:** 10.1101/2023.09.17.557749

**Authors:** Joshua S. Weinstock, Maya M. Arce, Jacob W. Freimer, Mineto Ota, Alexander Marson, Alexis Battle, Jonathan K. Pritchard

## Abstract

The effects of genetic variation on complex traits act mainly through changes in gene regulation. Although many genetic variants have been linked to target genes in *cis*, the trans-regulatory cascade mediating their effects remains largely uncharacterized. Mapping trans-regulators based on natural genetic variation, including eQTL mapping, has been challenging due to small effects. Experimental perturbation approaches offer a complementary and powerful approach to mapping trans-regulators. We used CRISPR knockouts of 84 genes in primary CD4+ T cells to perturb an immune cell gene network, targeting both inborn error of immunity (IEI) disease transcription factors (TFs) and background TFs matched in constraint and expression level, but without a known immune disease association. We developed a novel Bayesian structure learning method called Linear Latent Causal Bayes (LLCB) to estimate the gene regulatory network from perturbation data and observed 211 directed edges among the genes which could not be detected in existing CD4+ trans-eQTL data. We used LLCB to characterize the differences between the IEI and background TFs, finding that the gene groups were highly interconnected, but that IEI TFs were much more likely to regulate immune cell specific pathways and immune GWAS genes. We further characterized nine coherent gene programs based on downstream effects of the TFs and linked these modules to regulation of GWAS genes, finding that canonical JAK-STAT family members are regulated by *KMT2A*, a global epigenetic regulator. These analyses reveal the trans-regulatory cascade from upstream epigenetic regulator to intermediate TFs to downstream effector cytokines and elucidate the logic linking immune GWAS genes to key signaling pathways.

## Introduction

A primary mission of human genetics is to discover genetic variation that is associated with disease. Genome-wide association studies (GWAS) have identified thousands of variant-disease pairs in recent years, spanning disease, behavioral, and molecular phenotypes. Functional analyses of GWAS loci have revealed that most GWAS SNPs are non-coding, demonstrating that the effects of genetic variation on complex traits largely manifest through regulatory variation^1,2^. However, the identification of the molecular consequences of non-coding SNPs has proven challenging. Recent efforts have catalogued expression quantitative trait loci (eQTLs) across diverse tissues and contexts^3–6^. These eQTL studies have been very successful in identifying genetic variation that associates with expression variation in *cis*.

However, except for a small number of examples, the *trans*-regulatory cascade of these *cis*-acting genetic variants remains largely unknown. Recent analyses of the genetic architecture of complex traits have shown that the bulk (60-90%) of expression heritability is mediated through a constellation of *trans* effects which typically have small effects individually but have a large contribution in aggregate^7–9^. These *trans* effects are difficult to discover with natural genetic variation because their effect sizes are small and may only exist in contexts that are missed in bulk-tissue steady state models of gene expression^10–13^. Thus, alternative approaches are needed to map the trans-regulatory effects of *cis*-acting eQTLs.

We previously mapped the trans-regulators of three key autoimmune disease genes, *IL2RA, IL2*, and *CTLA4* in primary human CD4+ T-cells using CRISPR knock-outs (KOs)^14^. In contrast to natural genetic variation, experimental perturbations are unconstrained by natural selection, which enables the manipulation of gene expression in ways that are unlikely to be permitted by natural selection^15^. We therefore sought to apply this approach to inborn errors of immunity (IEI) genes, which are associated with monogenic immune disease spanning regulation and function^16^. Although hundreds of these genes have been reported, the transcriptional consequences of their loss of function remain largely uncharacterized. We selected 30 IEI transcription factors (TFs) for CRISPR ablation in human CD4+ T cells to both characterize their function and construct a regulatory network. CD4+ T cells have previously been implicated as a causal cell type in the pathology of many autoimmune traits, including rheumatoid arthritis, multiple sclerosis, type 1 diabetes, among others^17,18^. To enable characterization of the properties of the IEI TFs as a whole, we selected 30 background TFs that are matched to the IEI genes in terms of pLI^19^ and expression level in CD4+ T-cells but have not been implicated in GWAS of immune phenotypes. We also included 24 upstream regulators of *IL2RA* which we had previously perturbed using the same protocol^14^. In total, we perturbed 84 genes from three gene sets which we used to construct a high-fidelity gene network relevant to immune disease.

Building on recent advances in the causal inference literature^20,21^, we developed a novel statistical method for estimating causal GRNs from perturbation data. In contrast to differential expression or correlation analyses, incorporation of causal inference approaches enables the estimation of both direct and indirect regulatory effects, where edges are interpreted as direct effects. We emphasize that in this work the term ‘direct effect’ is used to convey that the effect of one gene on another is adjusted for confounding pathways among other perturbed genes, rather than a claim of physical interaction. Direct effects are useful because they facilitate a coherent interpretation of gene networks as directed probabilistic graphical models. Our approach differs from many other gene networks in two key ways: 1) because our network is derived from experimental perturbations, the edges are much more likely to be causal than the edges in a network estimated from observational co-expression data, where the constituent variation is often of an unknown genesis; 2) our method enables estimation of possibly cyclic graphs, rather than the common restriction to directed acyclic graphs (DAGs)^20,22–24^. Human genetics has identified several examples of cyclic regulatory behavior^25^, so the restriction of GRNs to DAGs represents an artificial constraint that we circumvent with appropriate statistical technology.

We report the causal, cyclic GRN derived from applying our novel statistical method to the 84 CRISPR KOs. Because this method is a Bayesian modification of the Linear Latent Causal (LLC) algorithm, we refer to our method as LLC Bayes (LLCB). Using our network, we systematically characterized the properties of genes that distinguish background TFs from IEI TFs and the *IL2RA* regulators. We show that although IEI TFs and *IL2RA* regulators are much more likely to have outgoing connections than background TFs, all the genes form a highly interconnected network, rather than distinct communities of disease and background genes. Across the entire network, we found that IEI and *IL2RA* regulators are more likely to disrupt immune specific signaling pathways than background TFs. We then identified nine coherent gene programs among the 84 KOs and their downstream genes, which we characterized using enrichment analyses to identify points of functional convergence in T cell biology. In addition to downstream characterization, we used GWAS summary statistic heritability analyses to estimate the contribution of gene program linked SNPs to immune trait heritability. This profiling highlighted the importance of a module comprised of key JAK-STAT-IL2 signaling regulators and *KMT2A*, a global epigenetic regulator that we observed to be upstream of classic IL2 signaling TFs and receptors, including *IRF4, STAT5B*, and *IL2RA*.

In summary, we perturbed a diverse set of genes to characterize the immune regulatory landscape and develop novel statistical methodology to characterize the CD4+ T cell network centered around immune disease genes. Our network reveals the entire trans-regulatory cascade of these gene programs and elucidates the transcriptional logic of immune GWAS loci.

## Results

### Perturbation of IEI TFs and matched background TFs

To construct a network enriched for genes relevant to immune disease in CD4+ T cells we perturbed 30 TFs from the IEI genes implicated in Mendelian forms of immune disease^16^. We also included 30 background TFs that were not annotated for immune function but were matched on gene constraint and expression to the IEI TFs in order to characterize the properties that distinguish IEI TFs. Lastly, to expand the breadth of our network, we integrated data from 24 previously mapped IL2RA regulators^14^. (Methods, Figure 1). We used CRISPR Cas9 ribonucleoproteins (RNPs) to perform arrayed perturbations in three donors as described in Freimer et al.^14^. We validated the efficiency of our CRISPR editing by genotyping the 60 additional perturbed samples, which indicated a high editing efficiency (Extended Data Figure 1A-B). Using bulk RNA-seq, we detected ∼13,000 genes that were expressed highly enough for analysis (Methods). As our data were generated in two batches, we performed stringent quality control of the RNA-seq data. We performed alignment and gene count quantification using one pipeline on the 84 samples and performed PCA analysis of the normalized expression data.Pathway enrichment analysis revealed that the first four PCs were associated with very broad biological phenomena including cell cycle regulation and ribosome activity. Because the PCs also captured batch effects, we included the first four PCs as covariates in downstream analyses. Regressing out PCs has previously been shown to improve inference of gene networks^26^.

**Figure 1.**
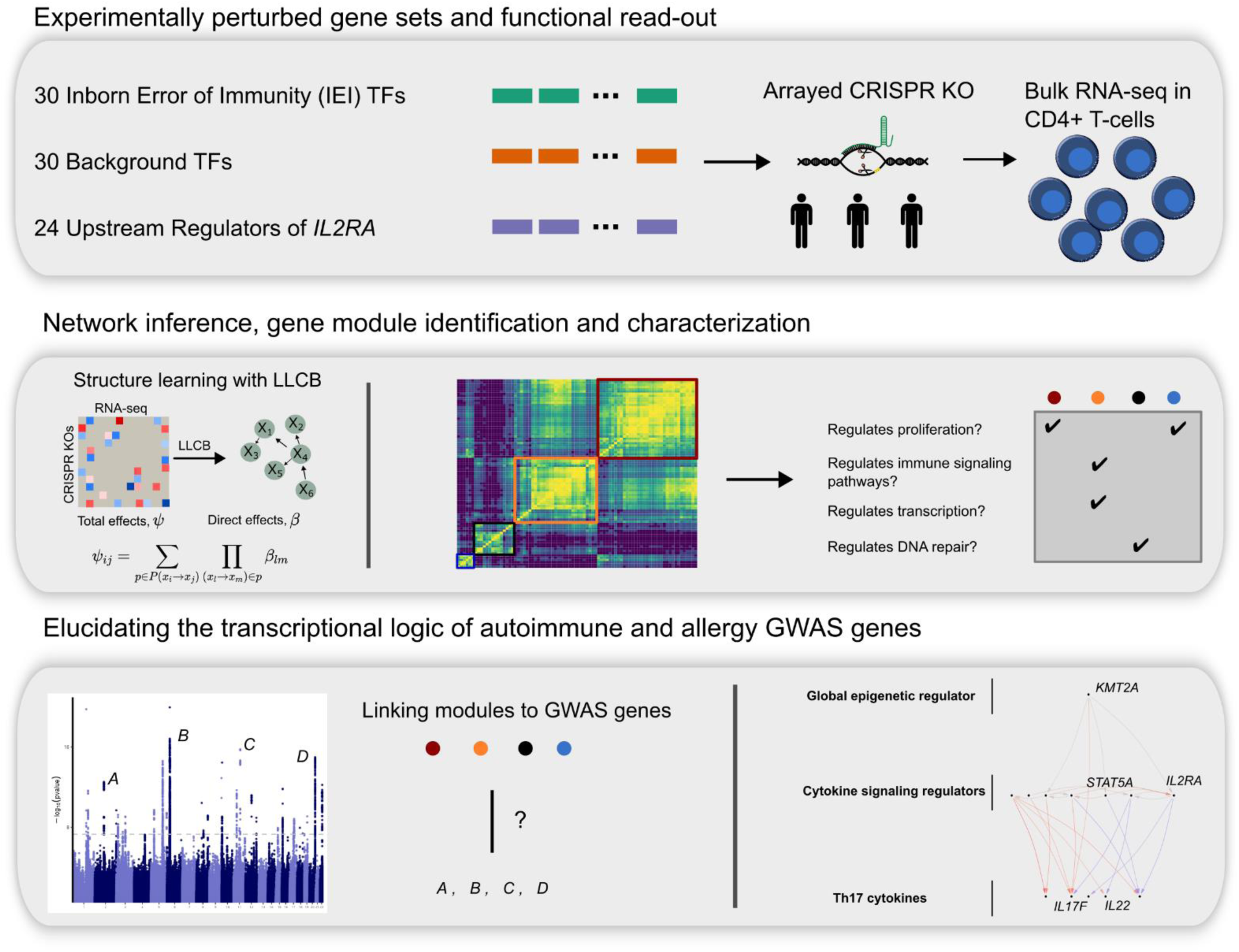
Study overview. Schematic describing the three gene sets that were perturbed with CRISPR knock outs and modeling of the gene network, network inference analyses and gene module identification, and integration with immune GWAS data.

Next, we developed a statistical method to estimate the GRN among the 84 genes. We extended the linear-latent-causal (LLC) method introduced by Hyttinen et al.^21^ by recasting the statistical estimand in a Bayesian framework, which enabled the incorporation of prior knowledge about the properties of biological networks. Briefly, LLC proceeds in three steps. First, the total effect ψ_*i,j*_ of a given perturbation of gene *X*_*i*_ on another gene, *X*_*j*_, is estimated on all observed (non-perturbed) genes. These total effects are estimated pairwise between all perturbed genes {*X*_*i*_: *i* ∈*J*} and all observed genes {*X*_*j*_: *j* ∈ *U*}. Second, a system of equations that relates ***ψ*** to the direct effects, ***β***, using trek rules is constructed. Third, this system of equations is solved to deconvolve ***ψ*** into ***β***. The conditions that permit identifiability of ***β*** for LLC include a collection of single gene perturbations among all nodes in the graph, which corresponds to our experimental design, indicating that we have a sufficient number of perturbations to identify ***β***. Because most of the 84 genes are TFs, the elements of ***β*** are likely to be greatly enriched for physical binding interaction and other mechanisms of direct transcriptional regulation. However, ***β*** may also capture post-transcriptional regulation mechanisms that manifest as statistical direct effects on expression. In this experiment we are unable to account for the effects of genes that were not perturbed, suggesting that some effects of unmeasured genes may be attributed to direct effects among the 84 perturbed genes.

We extended the LLC framework in two ways (Methods). First, we regressed out the first four expression PCs from the variance-stabilizing transform^27^ normalized expression data. Second, we estimated ***β*** in a Bayesian framework where we incorporated a graph prior, *π*(***β***). We included a penalty on the sum of the L1 norms of the columns of ***β***, which penalizes the number of incoming connections to a given gene. We included this penalty as it is known that the distribution of out-going connections from a gene has more dispersion than the distribution of in-coming connections. Following recent advances in differentiable DAG search^20,28,29^, we also included a gaussian prior over the norm of the spectral radius of ***β***, which enables indirect tuning of the degree to which ***β*** contains cycles. We performed inference using pathfinder, a recently developed approach to inferring posteriors using pseudo-Hessian optimizers applied to a variational inference objective^30^. We chose a variational inference approach rather than MCMC as MCMC approaches have been shown to be computationally very intensive when sampling over large discrete graph structures^23,24,31,32^. We termed this statistical method LLCB. We validated LLCB theoretically using simulations of cyclic GRNs (Methods).

### Network inference from LLCB reveals that the gene groups are highly interconnected

We then used LLCB to estimate the causal CD4+ GRN among the 84 genes (Figure 2A-B). We identified 350, 211, and 151 total edges (out of 6,972 possible) when thresholding |*β*_*ij*_| at 0.020, 0.025, and 0.030, respectively (Fig. 2A-C, Supplementary Table 1). We reported the network after thresholding on ***β*** because filtering on local-false sign rate (lfsr)^33^ resulted in very dense networks (67% network density at lfsr < 5 x 10^−3^), reflecting the challenges in estimation of uncertainty in graph structures.

**Figure 2.**
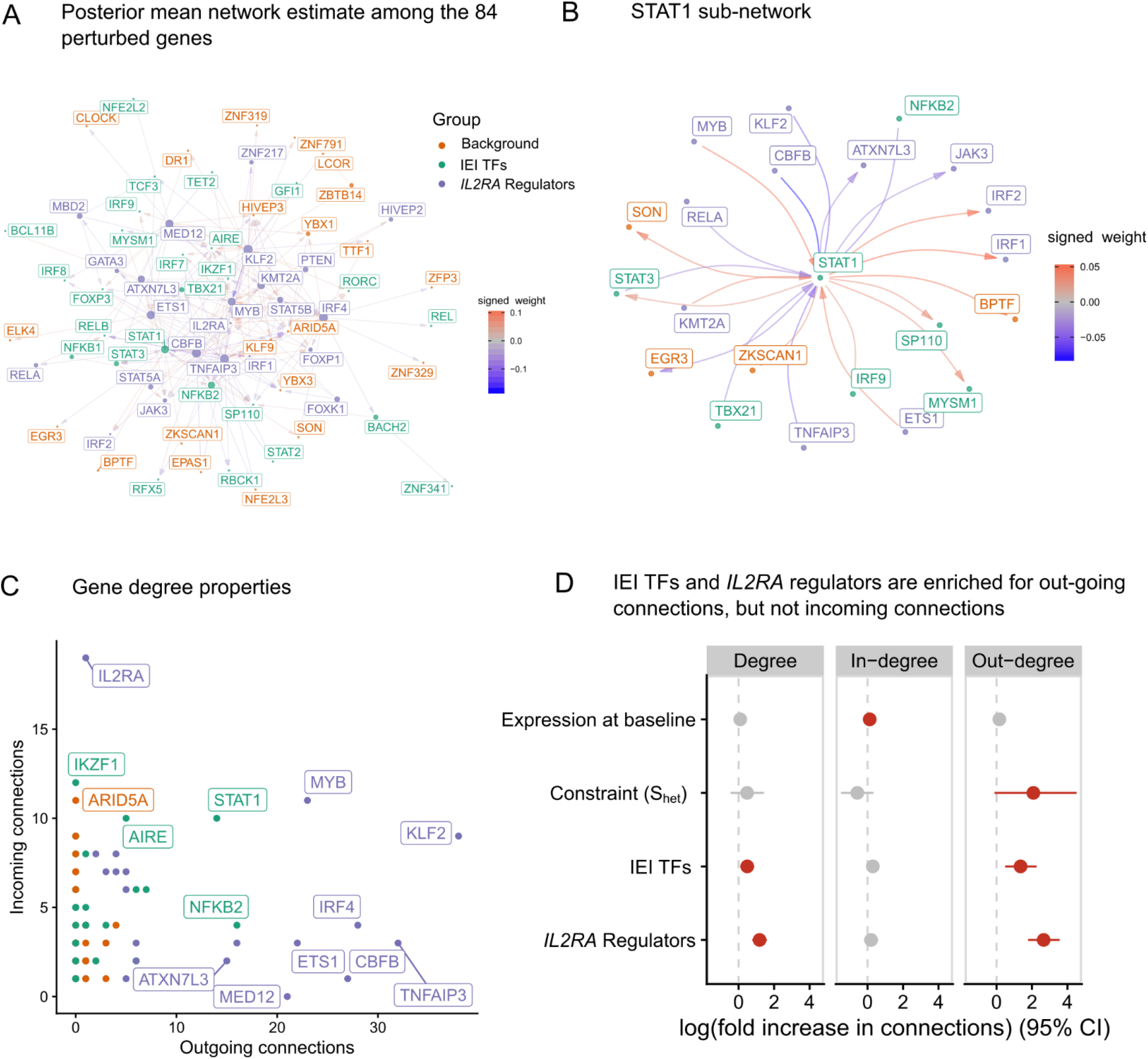
The gene network of the 84 perturbed genes. **A** Estimate of the directed network that describes how the 84 perturbed genes interact. The radius of each point is proportional to the degree of that gene. Arrows are used to indicate directionality of the edges, such that an arrow pointing into a gene indicates that it is being regulated by another gene. Positive values in the color scale indicate that the parent gene is a positive regulator of the child gene. **B** A sub-network centered around STAT1. **C** A scatterplot of the indegree and outdegree of each of the 84 genes. **D** Association analyses between gene properties and their in-degree, out-degree, and total degree.

To assess whether edges in this network estimate could be validated through orthogonal approaches, we compared our network estimate to two other estimates of the same network constructed from different sources. First, we constructed a GRN using ATAC-seq data that we previously generated for the 24 IL2RA regulators, permitting validation of a subset of the network. We gathered all possible enhancers of the 84 genes in CD4+ T cells using the predicted enhancer-gene pairs from the Activity-By-Contact^34^ model and cross-referenced the enhancer-gene pairs with differentially accessible chromatin (DAC) that we previously identified. We defined the children of a gene *i* based on those genes that had ABC enhancers that intersected with the DACs from the KO of gene *i*, and we refer to this network as the ABC-GRN (Supplementary Table 2). We observed a striking enrichment (∼4x) of edges in the LLCB estimate for the same edges in ABC-GRN, and this enrichment was robust to different |*β*_*ij*_| thresholds (Extended Data Figure 2). Second, we used an external estimate of the T-cell regulatory network reported in Green et al.^35^, which was estimated using curated pathway information and co-expression data. We similarly observed an enrichment of our edges in this external network (Extended Data Figure 3). Collectively, these two validations, derived from orthogonal data sources and modalities, show that our network estimate is replicable and reflective of biological properties.

We then asked whether the topological properties of genes distinguished the three gene groups. We computed the out-degree, in-degree, and total degree for each node, and we observed that the IEI TFs and *IL2RA* regulators were strongly enriched for out-going connections, and the control TFs were relatively depleted (Figure 2C). Consistent with the known properties of *IL2RA* as a receptor, as opposed to a TF, we observed many more direct incoming connections than direct outgoing connections. This result was likely facilitated by our inclusion of the downstream effectors of *IL2RA* signaling within the graph, such that downstream effects were more likely to be attributed to these genes, such as *STAT5A/B* and *JAK3*, rather than *IL2RA* itself. To identify the properties of genes that associated with their centrality in the graph, we performed negative-binomial regressions for three measures of node centrality, including gene group status, gene expression at baseline, and gene constraint as covariates. We defined gene constraint using a recently developed empirical Bayes estimator of *S*_*het*_, called GeneBayes^36^. *S*_*het*_ is defined as the degree of selection acting against heterozygous loss of function variants in a given gene and is more predictive of functional and clinical importance than related measures including pLI and LOEUF. We observed that even after adjusting for *S*_*het*_ and expression, *IL2RA* regulators and IEI TFs were strongly enriched for outgoing connections relative to control TFs but were not enriched for incoming connections (Figure 2D). Taken together, these data suggest that constraint is much more strongly associated with the number outgoing connections from a gene than the number of incoming connections, and that IEI regulators exhibit more outgoing connections than control genes, despite being matched for constraint.

We asked whether edges were enriched between genes that were members of the same gene group. To generate a null distribution, we permuted the edges of the network 2,000 times while preserving the gene degree distributions (Methods). Of the edges in the unpermuted network, 37% had the same parent and child node gene group. Of the permuted networks, 8% had more edges within groups than in the original (unpermuted) network, indicating that the three gene groups do not cluster distinctly in the unpermuted network (Extended Data Figure 4).

We then estimated indirect effects between pairs of genes, defined as the difference between the total effects and the direct effects *Δ*_*ij*_ = *ψ*_*ij*_ − *β*_*ij*_. The indirect effects can be interpreted as the sum of all effects of gene *i* on gene *j* that are not mediated through the direct effect *β*_*ij*_, and thus may include both proximal indirect effects comprised of short (< 3 genes involved) paths between the two genes or potentially distal effects from long, possibly cyclic paths. These indirect effects may include both instances of transcriptional regulation and post-transcriptional indirect effects. We observed that the bulk of variation in total effects (R^2^ = 99%) is explained by direct effects (Extended Data Figure 5), suggesting that direct effects between two genes are much larger than indirect effects. This observation is consistent with the intuition that indirect effects, which are defined as the product of several direct coefficients, are likely to be small unless all of the direct effects along the path are very large. Indeed, if all direct effects are less than 1.0 in magnitude, the product is guaranteed to be no larger than the smallest direct effect included in the path. We observed that the largest indirect effects were mediated by length-2 cycles with two large direct effects (Extended Data Figure 6). For example, we observed that *KLF2* and *MYB* regulate each-other in a length-2 negative feedback loop, which may help prevent aberrant proliferation.

### Trans-eQTL derived networks have limited overlap with the perturbation derived network

To compare our network estimate to one constructed from natural genetic variation, we first obtained the unfiltered trans-eQTL summary statistics from Yazar et al^6^, which contains the largest catalogue of CD4+ eQTLs mapped to date. We observed that only 24 of the 84 perturbed genes had at least one cis-eQTL (FDR < 0.01). The 24 genes with cis-eQTLs were much less constrained than the 60 without (difference in mean *S*_*het*_ = -0.07, 95% CI: (−0.15, 0.01)), corroborating our prior observations that eQTL discovery is biased towards genes tolerant to loss of function variation^15^. None of the 84 genes had a trans-eQTL, even at liberal significance thresholds (FDR < 0.30), indicating that this eQTL catalogue was incapable of recapitulating any of the edges in our GRN despite considering trans-eQTLs. To evaluate whether the absence of trans-eQTLs among the 84 genes was the result of trans-eQTL network sparsity, we tabulated the number of trans-eGenes in CD4 naïve and effector cells at FDR < 0.30, resulting in 12,185 trans-eGenes out of 16,025 tested genes. This implies that the probability of observing 84 randomly selected genes with no trans-eQTLs is 7 x 10^−53^, indicating that trans-eQTL sparsity alone cannot explain this observation. Collectively, these observations indicate that these TFs are strongly depleted of trans-eQTLs, potentially due to selective constraint, suggesting that mapping trans-regulators of highly constrained TFs with natural genetic variation is very under-powered at current sample sizes.

### Immune GWAS genes are enriched for regulation from IEI TFs and IL2RA Regulators

Next, we expanded our network analyses to include all 12,803 other genes that were expressed highly enough for analysis (Methods), which we refer to as non-perturbed genes. We estimated the effects of the 84 perturbed genes on the non-perturbed genes using two methods. First, we used a traditional differential expression approach using DESeq2^27^, where we regressed the normalized expression of each gene against a design matrix that included an indicator for the perturbation status of the sample, the donor identity, and the first four expression PCs. Next, we used mashr^37^ to perform statistical shrinkage of the differential expression estimates. We refer to these results as DEG-mashr estimates. To model the effects of multiple upstream TFs at the same time, we developed a novel statistical estimator of the bipartite graph (BG), which models the effects of the 84 perturbed genes on the 12,803 non-perturbed genes jointly in a single linear model. In contrast to a differential expression approach the BG model is less likely to detect redundant causal pathways. We term this approach the BG model (Figure 3A, Methods).

**Figure 3.**
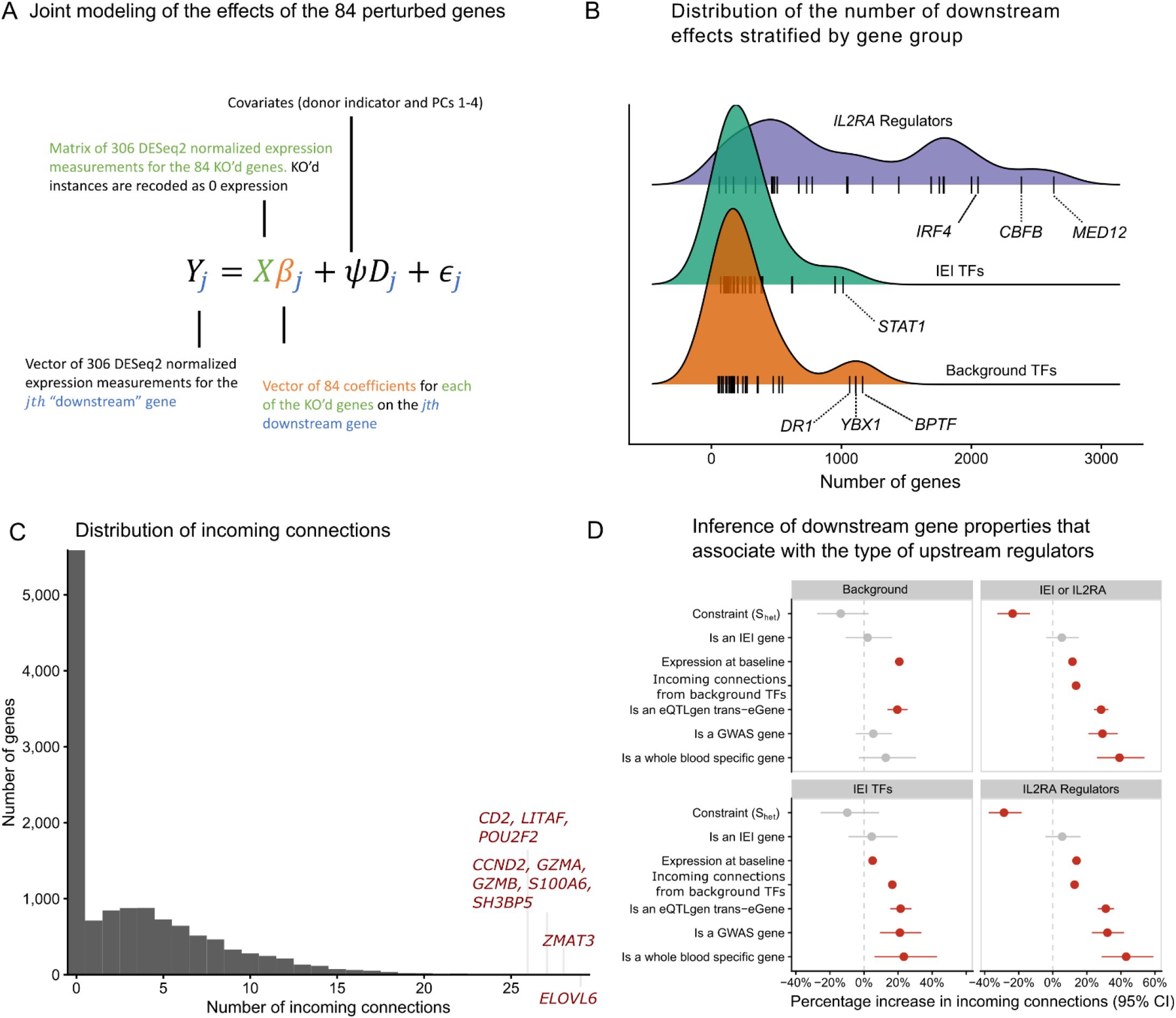
The landscape of downstream effects. **A** The statistical model used to relate the 84 perturbed genes to the expressed genes. **B** The distribution of the number of downstream effects for each of the 84 genes, stratified by gene group. Genes that are outliers with respect to their gene group distribution are labeled. **C** The distribution of indegree for each of the non-perturbed genes. Outlier genes are labeled. **D** Association between the properties of downstream genes and the gene-set of the upstream regulators. Coefficients are estimated with negative binomial regressions of the gene-set specific indegree.Downstream gene annotations are indicated on the y-axis and the facets are used to indicate the gene-set of the upstream regulator.

Among the non-perturbed genes, 7,299 (57%) had an incoming edge from at least one KO. Among the non-perturbed genes with at least one incoming edge, the median number of incoming edges was 5. The median number of downstream effects from the BG model was 251.5, ranging from 52 (*EGR3*) to 2,634 (*MED12*). Estimates from both the DEG-mashr and BG approaches (Supplementary Tables S3-5) revealed the striking enrichment of *IL2RA* regulators among the genes with the largest number of downstream connections (Fig. 3B). We observed that *MED12* and *CBFB* regulated more genes than any canonical T-cell transcription factor. *MED12* is a sub-unit of the mediator complex, which transmits signals from enhancer bound TFs to RNA-polymerase II bound at the promoter^38,39^. Despite its large effects, *MED12* has never been reported in any autoimmune GWAS, nor does it have a known cis-eQTL in CD4+ T-cells^6^, underscoring the value of perturbations for characterizing its function.

To our surprise, we also observed that three of the background TFs (*DR1, YBX1*, and *BPTF*) regulated more genes than any of the IEI TFs. The widespread effects of these three background TFs highlight the value of large-scale searches for upstream regulators, even in cell types with well annotated signaling pathways. Consistent with their large effects, these three TFs were highly constrained (*S*_*het*_ estimates of 0.38, 0.17, 0.30 for *DR1, YBX1, BPTF*). Although *BPTF* had no outgoing connections to the other 83 KO’d genes, it had an incoming connection from *STAT1*, suggesting that it may partially mediate the effect of *STAT1* on downstream genes. Among the 7,299 downstream genes with at least one incoming connection, there were 10 genes with at least 26 incoming connections (Fig. 3C), including genes involved in DNA damage response (*ZMAT3*), cell cycle regulation (*CCND2*), granzymes (*GZMA, GZMB*), and a T-cell cell costimulatory receptor (*CD2*).

Next, we asked which properties of the 12,803 non-perturbed genes were associated with regulation from the three gene groups. We performed a series of negative-binomial regressions of the incoming connections to non-perturbed genes, including six gene annotations as covariates (Figure 3D). We observed non-perturbed autoimmune GWAS genes were much more likely to be enriched for regulation from IEI TFs (∼20% enrichment) and *IL2RA* regulators (∼30% enrichment). *S*_*het*_ was negatively associated with incoming connections in three of the four regressions, consistent with our prior observation that gene constraint is more strongly associated with the number of outgoing connections from a gene than the number of incoming connections to the gene. We also observed that eQTL trans-eGenes were strongly enriched for incoming connections in each regression, suggesting that trans-eGenes reside in the periphery of the network with many incoming connections. Using GTEx, we also identified genes that were only expressed in whole blood and asked whether regulation of blood specific genes varied by the three gene groups. We observed that blood-specific genes were much more likely to be regulated by IEI TFs (∼20% enrichment) and *IL2RA* regulators (∼40% enrichment) than background TFs. Collectively, these observations highlight that although background TFs have similar graph centrality to IEI TFs, they are much less likely to disrupt cell type-specific transcriptional pathways.

### Gene modules link groups of genes to shared function

Next, we asked whether there were groups of the 84 perturbed genes with similar effects on downstream pathways among the 12,803 non-perturbed genes. Hierarchical clustering of the DEG-mashr results revealed the presence of nine gene modules (Figure 4), which we also grouped into a coarser set of super-modules. We remark that although the perturbed genes within each of these modules are mutually exclusive, the non-perturbed genes may overlap. To identify pathways that were regulated by these gene modules, we performed systematic enrichment analyses using KEGG genetic, signaling, and immune pathways^40^ (Figure 5A, Extended Data Figures 7-9, Supplementary Table S6).

**Figure 4.**
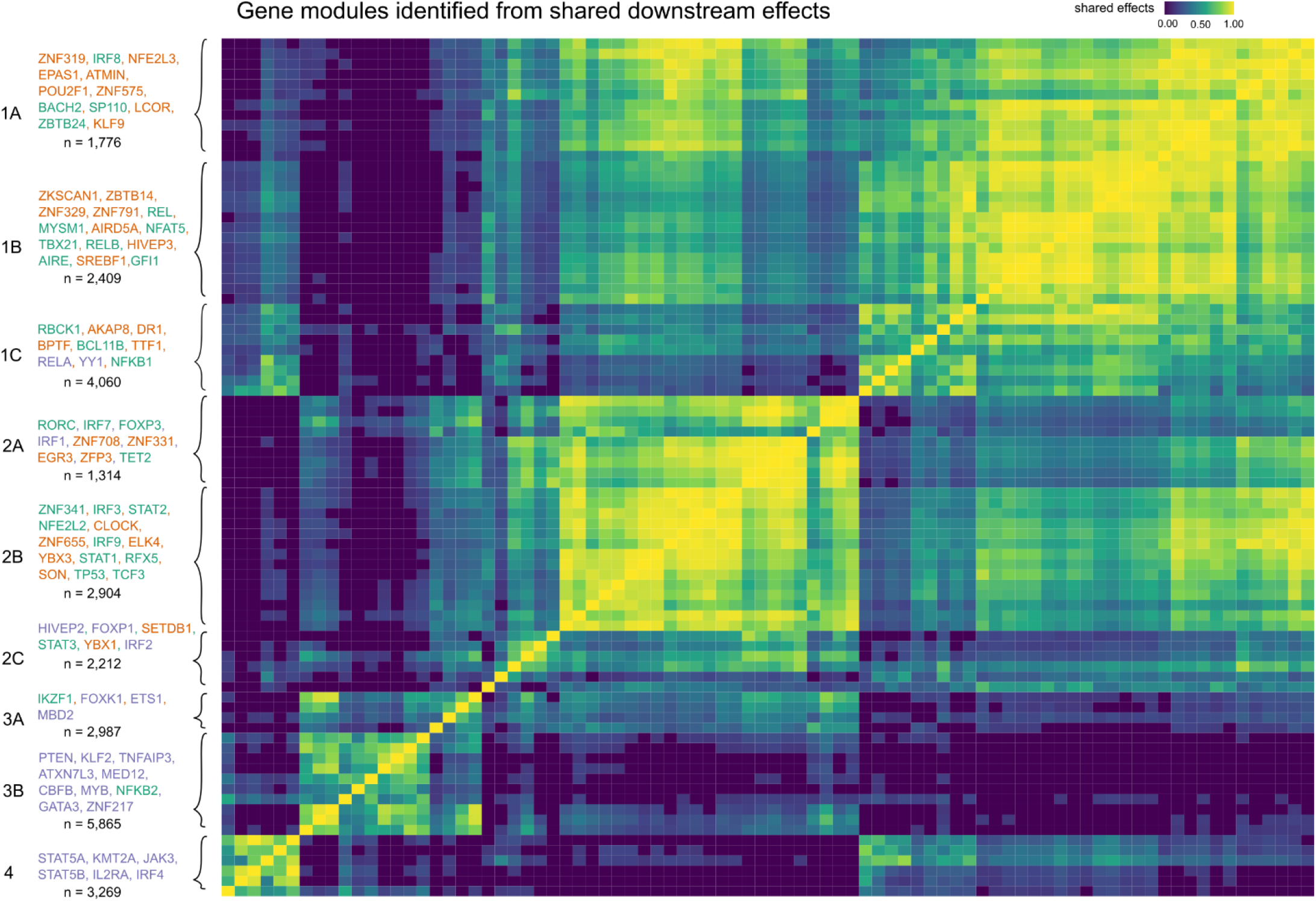
The discovery of gene modules. Hierarchical clustering is used to identify clusters of shared downstream effects. The upstream gene members within each module are labeled in the left-handed margin of the plot, and the gene group of each gene is indicated by the text color. The total number of genes in the module, including both upstream and downstream effects, is included under the list of genes.

**Figure 5.**
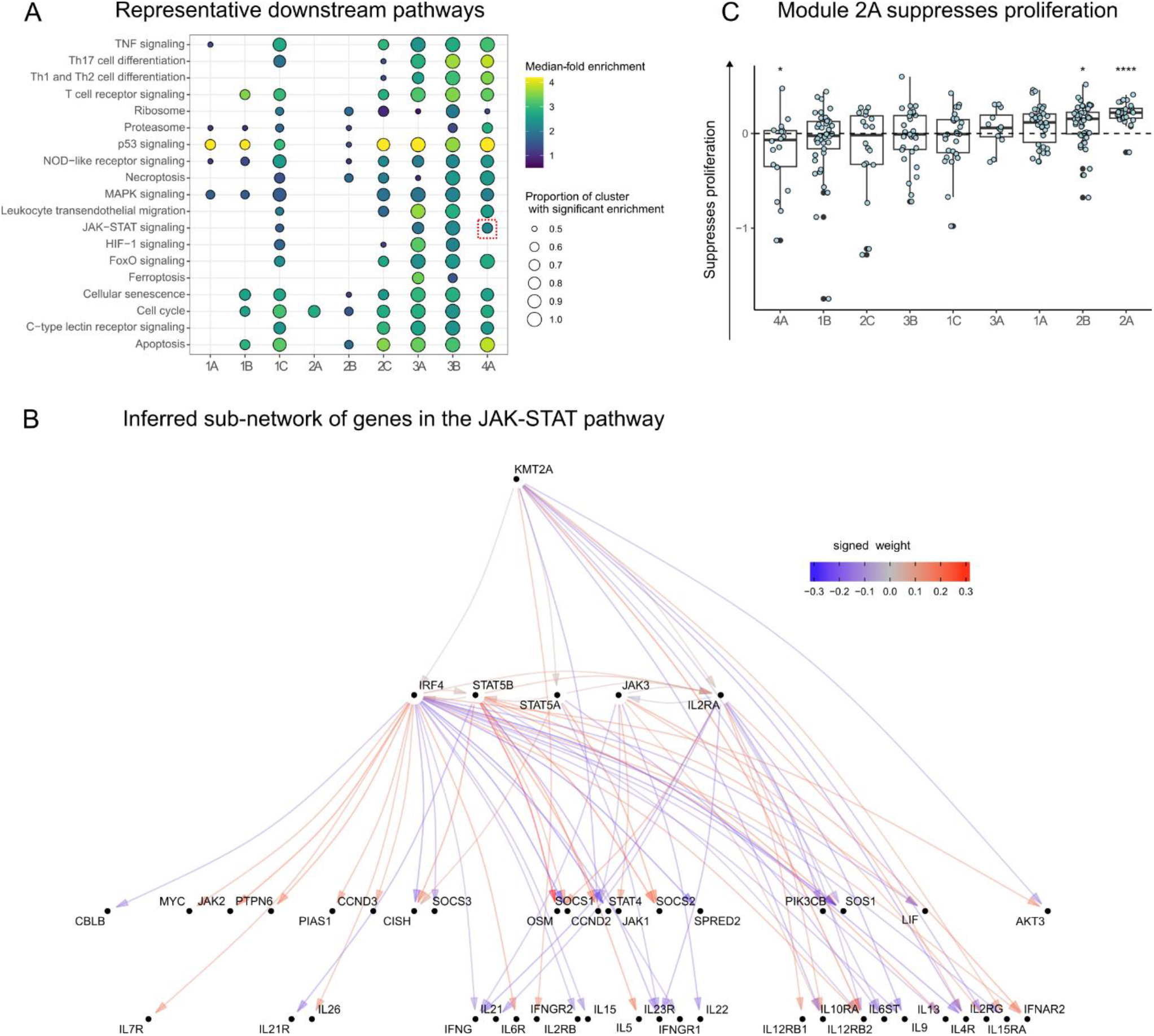
Gene module characterization. **A** Enrichment analyses of KEGG genetic, immune, and signaling pathways for each of the 84 perturbed genes, stratified by gene module. The JAK-STAT pathway is highlighted with a dashed-red box. **B** The JAK-STAT sub-network, which is organized such that cytokine genes are at the bottom and upstream regulators are at the top. **C** Effects of knock outs in the gene modules on a proliferation assay. Each point represents an individual gene perturbation sample plotted as the log2 fold change sample count as compared to AAVS1 KO control samples from the same donor. (*: p-value < 0.05, **** p-value < 0.001; n=3 donors per KO, the number of KOs per cluster is reflected in figure 4).

The perturbed genes in module 1 included 14 IEI TFs, 19 background TFs, and two *IL2RA* regulators (*RELA* and *YY1*). The perturbed genes in modules 1-2 were primarily IEI and background TFs, and modules 3-4 were primarily *IL2RA* regulators. We observed that module 1A was enriched for disruption of MAPK and p53 signaling. Module 1B included T-bet (*TBX21*), a transcription factor that is required for interferon-gamma production and the Th1 phenotype^41^, and three members of the Rel family (*NFAT5, RELB*, and *REL*), sub-units of NF-κb, a transcription factor complex that plays a role in T-cell activation^42^. Surprisingly, this cluster also included four background TFs without any annotated immune function (*ZNF329, ZNF791, ZBTB14*, and *ZKSCAN1*). ZBTB7B has been observed to be required for CD4+ commitment, and interacts with NF-κB^43^, but many other members of the ZBTB family, including *ZBTB14*, remain relatively uncharacterized. The high proportion of shared effects between *ZBTB14, T-bet*, and the Rel family proteins suggests that *ZBTB14* may have similar function to *ZBTB7B*.

Genes in super-module 2 were enriched for effects on cell cycle regulation and apoptosis. Modules 3-4 were much more strongly enriched for *IL2RA* regulators than clusters 1-2. Consistent with their annotation, every gene in module 3-4 had downstream effects on the JAK-STAT and chemokine signaling pathways. Surprisingly, *KMT2A*, a methylation writer clustered in the same module as *JAK3, STAT5A, STAT5B, IRF4*, and *IL2RA*. Although translocations of *KMT2A* have been shown to cause lymphoid malignancy^44^, it has no annotated function in non-mutated cells in the JAK-STAT pathway^45^. We then examined the structure of module 4 (Figure 5B), observing that *KMT2A* is upstream of *IRF4, STAT5A*, and *IL2RA*, and directly regulates several downstream effector cytokines through pathways not mediated by the other perturbed genes.

Several modules were strongly enriched for cell cycle and proliferation pathways. To determine if there was a uniform effect on *in vitro* expansion within any of the modules, we quantified the number of live cells per KO compared to cells where the guide RNA targeted the safe harbor locus AAVS1 from the respective donor. All members of module 2A, which was enriched for cell cycle effects, showed a mean increase in cell counts across three donors as the result of the perturbation. Collectively, the module had a 1.16-fold increase in live cells when KO’d compared to the controls, suggesting that genes in 2A function as proliferation repressors (Figure 5C). Concordant with these observations, a recent report described the proliferation promoting effects of disruption of a module 2A member, *TET2*, in CAR-T cells^46^. Our analyses suggest that other members of 2A may have similar properties to *TET2* and thus may represent a group of genes that could be perturbed to alter engineered T-cell function. Several upstream members of 2A upregulated three of four CDKN genes which inhibit cyclin dependent kinases and potentially lead to reduced cycling (Extended Data Figure 10). Taken together, our inference of gene modules recapitulates known regulators of immune signaling pathways and identifies novel members of these modules.

### Heritability analyses link gene modules to immune disease risk

We then asked whether SNPs that were linked to the nine gene modules were enriched for heritability of autoimmune traits. We included GWAS summary statistics for 10 phenotypes from a combination of Finngen and disease specific consortia^47,48^. After linking SNPs to each of the nine modules using the Activity-By-Contact method^34^, we used LD score regression^2,49^ to estimate the contribution of these SNPs to the heritability to eight autoimmune traits and two allergy traits (Methods, Supplementary Table S7). As a reference point, we also included a group of SNPs linked to genes that were not regulated by any of the 84 genes, which we term module 0. To adjust for confounding genomic annotations, we included the LD-score baseline model. We observed that module 4 SNPs were potent contributors generally, as half of the traits analyzed were enriched (Figure 6A, Extended Data Figure 11, Supplementary Table S8). Across the traits, there was substantial heterogeneity in the effects of modules. For example, only 4A and 2B SNPs were associated with psoriasis heritability, while 1A, 2B, 3A, 3B, and 4A all contributed to rheumatoid arthritis heritability. Among the baseline module 0, only multiple sclerosis was enriched. Remarkably, module 1B contributed little to heritability enrichment of any trait despite including TBX21.

**Figure 6.**
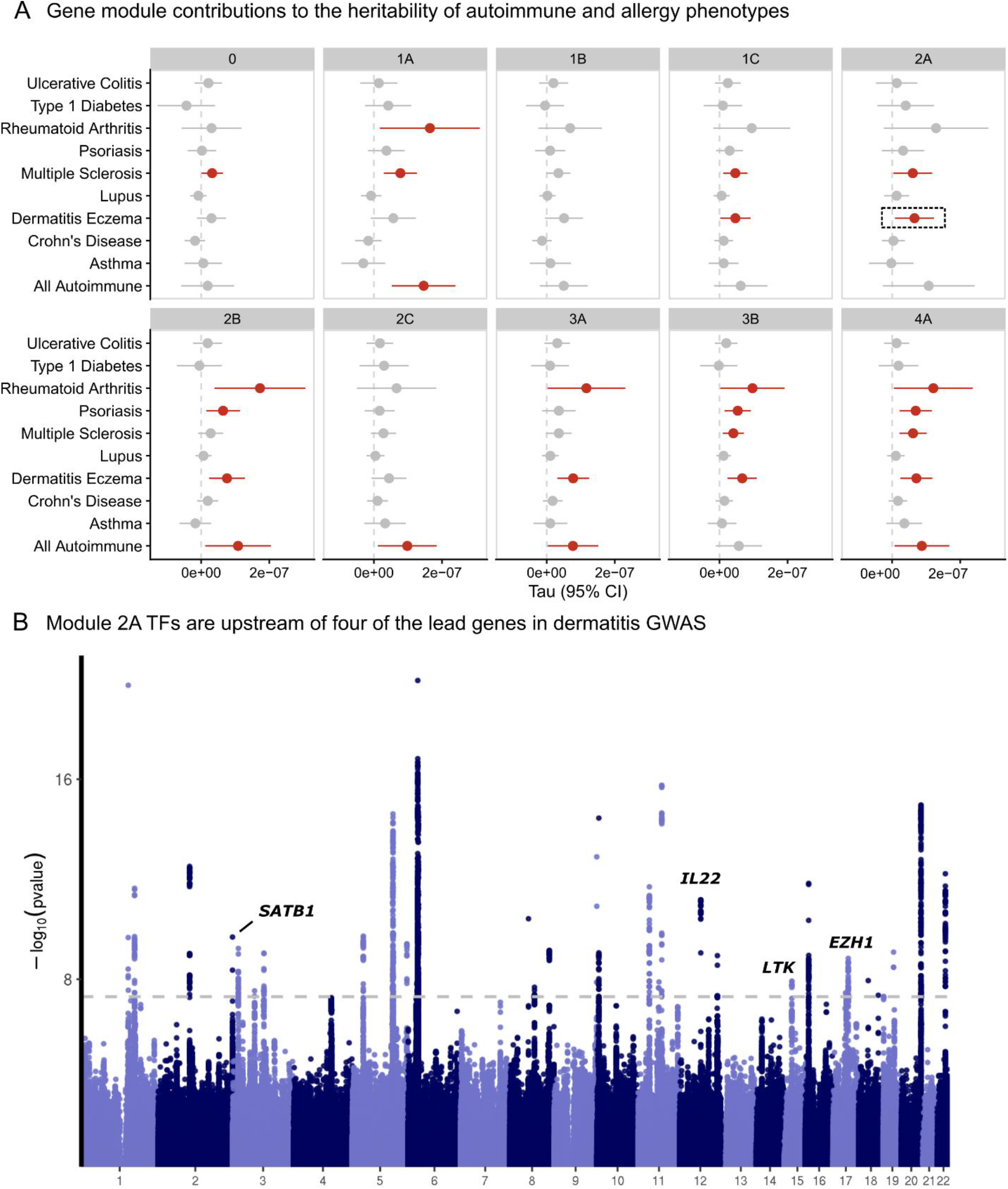
Contribution of SNPs linked to the gene modules to heritability of autoimmune and allergy phenotypes. **A**. Estimated *τ* coefficients from LD score regression are plotted for each gene module and phenotype. Module 0 is defined as genes that were not included in any module but are still expressed in CD4+ T cells. **B**. Exemplar analysis annotating the fine-mapped genes from a Finngen dermatitis GWAS based on their presence in module 2A.

We also observed that module 2A, which was strongly enriched for effects on cell cycle regulation pathways, was enriched for regulation of atopic dermatitis GWAS genes. Next, we annotated the fine-mapped signals from the dermatitis GWAS. Of the 44 credible sets, 34 were linked to genes. Of these 34 hits, four were regulated by the module 2A TFs, including *SATB1, IL22, LTK*, and *EZH1* (Figure 6B). Given the putative effects of module 2A on cell proliferation, we then cross-referenced these four genes with cell proliferation annotation pathways. *LTK* is a receptor with tyrosine kinase activity and may contribute to proliferation through activation of the PI3K signaling pathway^50^. Similarly, *IL22* has also been reported to regulate PI3K signaling^51^. Taken together, these analyses highlight the value of unbiased module discovery for identifying specific pathways that contribute to trait heritability. We illustrate how module 2A TFs regulate a subset of dermatitis GWAS genes that have been implicated in PI3K signaling, a common proliferation pathway.

### The transcriptional logic linking the JAK-STAT module to immune GWAS genes

Given the substantial contribution of module 4 to autoimmune and allergy phenotype heritability and its large effects on T cell differentiation, we integrated multiple functional assays to elucidate the fine-grained structure of module 4. We observed that *KMT2A* was a positive regulator of IL17F and IL21 expression, two Th17 secreted factors (Figure 7A). We also observed concordant decreases in chromatin accessibility near (5.7 kb and 40 kb upstream of TSS) IL17F and IL21 upon KO of *KMT2A*. Notably, IL17F had a striking decrease in expression (−5.9 log2 fold change) in the *KMT2A* KO. We then intersected the differentially accessible chromatin regions from the *KMT2A* KO condition with each of the KOs within module 4 and observed that *STAT5B* shared several differentially expressed sites, including regions upstream of IL17F (Extended Data Figure 12). An additional Th17 secreted factor, IL22, also had a shared region between the two conditions, although the transcript was only differentially expressed in the *STAT5B* KO. The *STAT5B* KO also abrogated chromatin accessibility 5.7 Kb upstream the IL17F promoter. Concordant with these observations, ChIP-seq data generated in IL-2 stimulated CD4+ T cells confirmed direct binding of *STAT5B* (Figure 7B). Because *KMT2A* is a methyltransferase that deposits activating methylation marks on H3K4, we then asked whether H3K4me3 was present in these same peaks in Th17s stimulated with IL-2, finding that H3K4me3 marks were indeed present in the differentially accessible peaks (Figure 7B). These observations led to us suggest the following mechanism for the regulatory logic of module 4: *KMT2A*, a global epigenetic regulator of transcription, collaborates with downstream factors, including *STAT5B*, to positively regulate IL17F through modulation of activating histone marks and chromatin remodeling of a regulatory element that is likely an IL17F specific enhancer in Th17 cells.

**Figure 7.**
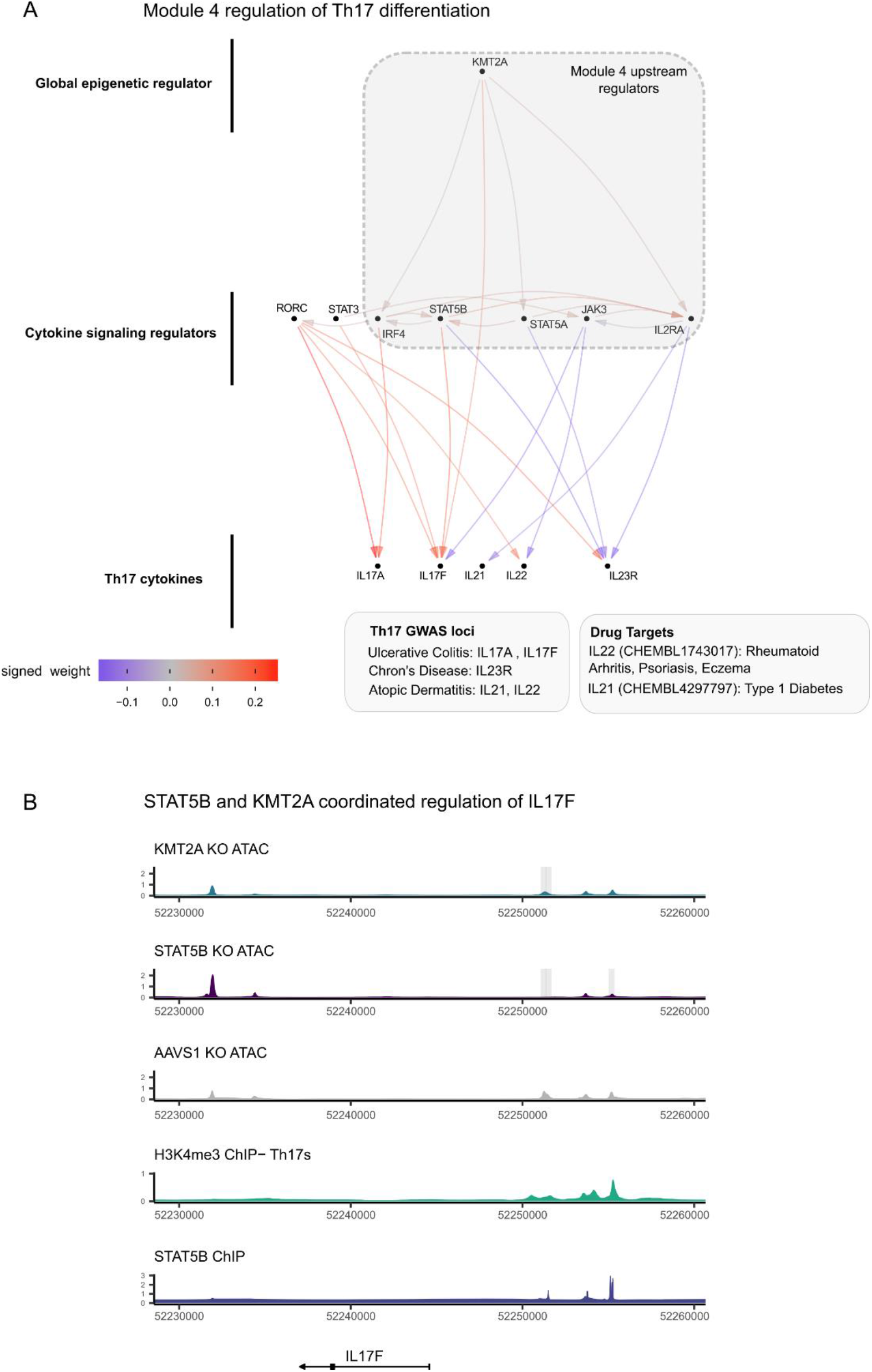
The transcriptional logic linking module 4 to GWAS loci. **A**, The sub-network of module 4 and Th17 cytokines. **B**, locus plot including tracks describing the functional characteristics of the region. Each track is constructed from publicly available ChIPseq data (methods) or ATAC-seq data from Freimer et al. Grey boxes indicate significantly different regions between the respective KO and AAVS1 control KO ATAC data (padj < 0.05, n = 3 donors per KO). The Y-axis displays normalized counts.

These observations suggest that cis-regulatory elements near *KMT2A* may harbor autoimmune risk variants. To assess this hypothesis, we examined recent biobank GWAS in UKB^52,53^, Finngen^47^, and Biobank Japan^54^ (BBJ) for variants associated with autoimmune phenotypes near *KMT2A*. The A-allele of rs45480496, a common variant (MAF of 21% in TOPMed^55^) 36Kb from the TSS of *KMT2A*, is suggestively associated with autoimmune disease (“diseases marked as autoimmune origin”, OR = 1.04, pvalue = 2 x 10^−7^) in Finngen and was also reported as suggestive hit in a BBJ-UKB meta-analysis^56^ (“autoimmune multi-trait”, OR = 1.08, pvalue = 2 x 10^−6^). A meta-analysis of these two signals results in genome-wide significance (pvalue = 2 x 10^−12^, Extended Data Figure 13) for this variant. We then looked for functional evidence linking rs45480496 to *KMT2A*. Although rs45480496 has not yet been reported as an eQTL for *KMT2A*, lookup of the SNP in a promoter Hi-C capture in immune cells^57^ revealed that it resides in a regulatory element that interacts with the promoter of *KMT2A* in megakaryocytes, naïve CD4s and CD8s, and effector CD4s and CD8s. Concordant with these observations, lookup of rs45480496 in regulomedb^58^ indicated that it is in an active enhancer in Th17 cells. The haplotype that rs45480496 tags also intersects with a predicted *KMT2A* enhancer in CD4+ T-cells from the ABC model^34^. Although the variant to gene predictions from OpenTargets^59^ suggest that other causal genes are possible in this locus, we remark that these predictions are made without knowledge of the causal cell type for a given phenotype. Collectively, these data report a novel risk locus for autoimmune traits upstream of *KMT2A* which likely contains a *KMT2A* enhancer.

## Discussion

Human genetics has been remarkably productive in discovering complex-trait associated SNPs.

There are now several resources to map the effects of these SNPs to molecular phenotypes in *cis*, however, the development of maps of the regulatory cascades of these SNPs has progressed much more slowly. Enabled by recent innovations in large-scale perturbation technologies, we are now able to systematically perturb large numbers of genes in primary human cell contexts. These perturbations complement natural genetic variation approaches to mapping trans-regulators as they facilitate the examination of biological variance that is unlikely to be observed in healthy cells. After network inference with LLCB, we observed 211 trans-regulatory causal connections in our upstream GRN, none of which were reported in the largest catalogue of CD4+ eQTLs performed to date^6^.

We developed LLCB to infer the gene network which builds upon recent advances in the structure learning literature to estimate a graph with edge weights that are interpretable as direct effects. This stands in contrast to the majority of effect estimates reported in the functional genomics literature, which primarily report estimates from differential expression analyses performed separately in each perturbed gene. These estimates confer results that are difficult to interpret because they do not attempt to adjust for confounding pathways in the GRN, which are known to be highly abundant in biological networks. We use LLCB to estimate the topology and effect size of these confounding pathways. We found that direct effects were generally much larger than indirect effects in magnitude, and that the largest indirect effects were mediated by local feedback cycles.

Using experimental perturbations, we investigated the properties of IEI TFs which are infrequently mutated in natural genetic variation. We performed a series of systematic analyses that delineate the commonalities and differences among the IEI TFs, background TFs, and *IL2RA* regulators. Consistent with our previous report^14^, we found that the *IL2RA* regulators were potent regulators of downstream effects. Both the IEI TFs and *IL2RA* regulators were enriched for being upstream and were much more likely than background TFs to disrupt autoimmune GWAS loci and whole blood specific genes even after adjustment for gene constraint. We also observed that the topology of the regulatory network is strongly associated with selective constraint. *S*_*het*_ was among the best predictors of the topological properties of the perturbed genes: *S*_*het*_ was strongly associated with the number of outgoing connections of a gene, but not the number of incoming connections. This is reflected in the dense downstream network identified for the *IL2RA* regulators with overall high levels of constraint, compared to the other TF groups. Overall, the difference in enrichment based on *S*_*het*_ suggests that the centrality of genes is best expressed as a multi-dimensional construct. This further highlights the value of estimating GRNs with directed edges, as opposed to estimating undirected graphs from observational co-expression data, as the richer graphical structure enables much more granular topological analyses.

Utilizing the novel connections in the GRN, we report several observations that improve annotation of canonical immune pathways. We observed that three of the background TFs (*DR1, BPTF*, and *YBX1*) regulated more downstream genes than any of the 30 IEI TFs, including *TBX21*, a master regulator of Th1 differentiation. After identifying gene modules and their downstream pathways, we observed multiple novel members of canonical gene modules, including *KMT2A* in the JAK-STAT pathway. We observed that *KMT2A*, a methyltransferase that deposits activating methylation marks, modulated the expression of canonical IL-2 signaling TFs. *KMT2A* collaborated with these TFs to upregulate IL17F, a pro-inflammatory cytokine that is secreted by Th17 cells, indicating that *KMT2A* is an under-appreciated regulator of the IL2-JAK-STAT axis and Th17 activation. Meta-analysis of biobank autoimmune GWAS revealed a novel risk locus in a Th17 enhancer upstream of *KMT2A*. Collectively, these observations suggest that *KMT2A* inhibitors may be a productive therapeutic avenue for autoimmune disease.

Although we have demonstrated that our regulatory network is useful for discovery of novel immune pathway biology and that it is validated by orthogonal data modalities, our study is not without limitations. The perturbation of additional genes in more donors, cell types, and cell contexts would undoubtedly result in increased discovery. The restriction to transcriptional regulation also inhibits the interrogation of post-translational regulation, which makes the interpretation of edges from genes where post-translational regulation important challenging. This suggests that the STAT proteins, which are known to be sensitive to phosphorylation, may regulate more genes than is estimated in our transcriptional network. The use of a bulk expression read-out, although more sensitive to genes with low expression than single cell assays, also precludes the analysis of more granular cell types and contexts.

In conclusion, we describe the gene regulatory network of key CD4+ T cell regulators. This network enabled both the broad characterization of the properties of immune disease genes and the discovery of novel regulatory connections between TFs and signaling pathways that modulate immune disease genes. We anticipate that our approach can be applied in other cell types and contexts to generate maps of the molecular consequences of regulatory variation of disease genes.

## Methods

### Cell Isolation and expansion

Primary CD25-CD4+ effector T cells were isolated from fresh Human Peripheral Blood Leukopaks (STEMCELL Technologies, #70500) from healthy donors, after institutional review board–approved informed written consent (STEMCELL Technologies). Peripheral blood mononuclear cells (PBMCs) were washed twice with a 1X volume of EasySep buffer (DPBS, 2% fetal Bovine Serum (FBS), 1mM pH 8.0 EDTA). The washed PBMCs were resuspended at 200E6 cells/mL in EasySep buffer and isolated with the EasySep™ Human CD4+CD127lowCD25+ Regulatory T Cell Isolation Kit (STEMCELL Technologies, #18063), according to the manufacturer’s protocol. Cells were seeded at 1x10^6^ cells/mL in complete RPMI-1640 supplemented with 10% FCS, 2 mM L-Glutamine (Fisher Scientific #25030081), 10 mM HEPES (Sigma, #H0887-100ML), 1X MEM Non-essential Amino Acids (Fisher, #11140050), 1 mM Sodium Pyruvate (Fisher Scientific #11360070), 100 U/mL Penicillin-Streptomycin (Sigma, #P4333-100ML), and

50 U/mL IL-2 (Amerisource Bergen, #10101641) and stimulated with 6.25 uL/mL ImmunoCult™ Human CD3/CD28/CD2 T Cell Activator (STEMCELL Technologies, #10990). Following activation and electroporation, cells were split 1:2 every other day to maintain an approximate density of 1x10^6^ cells/mL.

### Cas9 RNP preparation and delivery

Custom crRNAs (Dharmacon) and Dharmacon Edit-R CRISPR-Cas9 Synthetic tracrRNA (Dharmacon, #U-002005-20) were resuspended in Nuclease Free Duplex Buffer (IDT, #11-01-03-01) at 160uM stock concentration. In a 96 well plate, each crRNA was combined with tracrRNA at a 1:1 molar ratio and incubated at 37°C for 30 minutes. Single-stranded donor oligonucleotides (ssODN; sequence: TTAGCTCTGTTTACGTCCCAGCGGGCATGAGAGTAACAAGAGGGTGTGGTAATATTACGGTACCGAGCACTATCGATACAATATGTGTCATACGGACACG, 100uM stock) was added to the complex at a 1:1 molar ratio and incubated at 37°C for 5 minutes. Finally, Cas9 protein (MacroLab, Berkeley, 40 μM stock) was added at a 1:2 molar ratio and incubated at 37°C for 15 minutes. The resulting RNPs were frozen at -80°C until the day of electroporation. 48 hours following effector T cell activation, the cells were pelleted at 100x g for 10 minutes and resuspended in room temperature Lonza P3 buffer (Lonza, catalog no. V4XP-3032) at 1.5x10^6^ cells per 20 ul P3. The cells were combined with 5 ul aliquots of the thawed RNPs, transferred to a 96-well electroporation cuvette plate (Lonza, #VVPA-1002) and nucleofected with pulse code EH-115. Immediately following electroporation, the cells were gently resuspended in 90 ul warmed complete RPMI with IL-2 and incubated at 37 C for 15 minutes. After recovery, the cells were cultured in 96 well plates at 1x10^6^ cells/mL for the duration of the experiment. To prevent edge effects, the guides were randomly distributed across each plate and the first and last column of each plate was excluded, being filled instead with PBS to prevent evaporation.

### RNA isolation and library preparation

8 days after T cell isolation and activation, the cells were pelleted and resuspended at 1x10^6^ cells per 300 ul of RNA lysis buffer (Zymo, #R1060-1-100). Cells were pipette mixed and frozen at -80 until RNA isolation was performed. RNA was isolated using the Zymo-Quick RNA micro prep kit (#R1051) according to the manufacturer’s protocol with the following modifications: After thawing the samples, each tube was vortexed vigorously to ensure complete lysis prior to loading into the extraction columns. In lieu of the kit provided DNAse, RNA was eluted from the isolation column after the recommended washes and digested with Turbo-DNAse (Fisher Scientific, AM2238) at 37 C for 20 minutes. Following digestion, RNA was purified using the RNA Clean & Concentrator-5 kit (Zymo, #R1016) according to the manufacturer’s protocol. The resulting purified RNA was submitted to the UC Davis DNA Technologies and Expression Analysis Core to generate 3′ Tag-seq libraries with unique molecular indices (UMIs).

Barcoded sequencing libraries were prepared using the QuantSeq FWD kit (Lexogen) for multiplexed sequencing on an Hiseq 4000 (Illumina).

### Cell proliferation quantification

One replica plate of cells from each donor was run on the Attune NxT Flow Cytometer (Thermo Fisher) within 24 hours of cell lysis for RNA extraction. Cell density per knock-out condition was quantified by the Attune using an equi-volume amount of sample. Counts were normalized to the mean AAVS1 density for the respective donor.

### RNA-seq alignment and gene count quantification

Adapters were trimmed from fastq files with cutadapt^60^. Low-quality bases from reads were trimmed using the Phred algorithm implemented in seqtk^61^. Reads were then aligned with STAR^62^ and mapped to GRCh38. Gene counts from deduplicated reads were quantified using featureCounts^63^. Sample quality control reports were generated with Fastqc^64^, rseqc^65^, and Multiqc^66^.

### Gene filtering and PCA analysis

Genes were first filtered to those with at least 10 counts in at least five samples. PCA was then performed on the variance stabilizing transformed^27^ (vst) counts of the 500 most variable genes. Three outlier samples were excluded and then the above process was repeated. The PCs were then assessed for association with batch effects and very broad cellular pathways. PCs 1-2 associated with batch effects, and PCs 3-4 were associated with cell cycle state, suggesting that PCs 1-4 should be included as covariates or otherwise adjusted for in downstream analysis.

### Differential expression analysis

Differential expression analysis was performed using DESeq2^27^, including donor identity, PCs 1-4, and the KO as predictors of the response. Donor identity and PCs 1-4 were included as covariates to mitigate their confounding effects on gene expression. We emphasize that the statistical estimand in this analysis the total effect of the perturbation of a given gene on the readout gene. This effect may include several indirect paths between the perturbed gene and the readout gene.

### LLCB

We formulate the GRN as a graph ***G*** = (***X, β***), where the *P* nodes ***X***_*1*_, …, ***X***_*p*_ are each a vector of the vst normalized gene expression values. We restrict this analysis to the 84 KO’d genes reflecting the importance of satisfying the identifiability condition described in Hyttinen et al. ***β*** is the adjacency matrix describing the direct linear effects between genes, where the rows encode the parent genes and the columns encode the children genes. We then construct a covariate matrix ***W*** where the columns ***W***_1_, … ***W***_*l*_ indicate *l* covariates to regress out. We then orthogonalize ***X*** based on ***W*** with the transformation 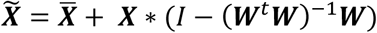, where 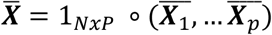. We add back in the column means 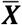to roughly preserve the original scale of ***X***. In ***W***, we include the donor identity and first four PCs as covariates.

We define *KO*_*j*_ as the indices indicating the samples in which ***X***_*j*_ was intervened upon and we set *O*_*j*_ = {1, …, *N*} *− KO*_*j*_. We define *C* as the indices in which safe-harbor AAVS1 control samples were used. For all *j* = 1, …, *P* we recode 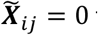 for all *i* ∈ *KO*_*j*_. This reflects our belief that the CRISPR KOs were effectively forcing the normalized functional transcript abundance to 0, i.e., we assume perfect interventions.

We then estimate ***β*** in two steps: 1. Estimate the total effects ψ_*ij*_ between every pair of genes (*i, j*) ∈ *P x P*; 2. Estimate ***β*** from ***ψ*** using a modification of the LLC algorithm^21^. To estimate ψ_*ij*_, we first center and scale 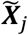based on its mean and standard deviation in the control samples *C*. Then, we use OLS to estimate the total effect of 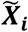 on 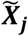, limiting the samples used to {*KO*_*i*_, *C*}, such that we exclude all instances in which the child node 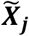 has been KO’d. This analysis results in the matrix of estimated total effects, 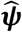. We emphasize that these coefficients are on a correlation scale because of the standardization procedure.

We assume asymptotic stability^21^ over the true β, which is equivalent to assuming that the largest eigenvalue is less than 1. Because we know β is asymptotically stable, the following decomposition of true effects into direct effects is coherent:

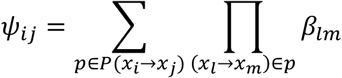

This relationship indicates that total effect of a gene *i* on gene *j* is the sum of all possible paths between them, where the value of an individual path is defined by the product of direct effects along that path.

To estimate β from 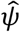, we use the LLC procedure Algorithm 1:

#### Algorithm 1

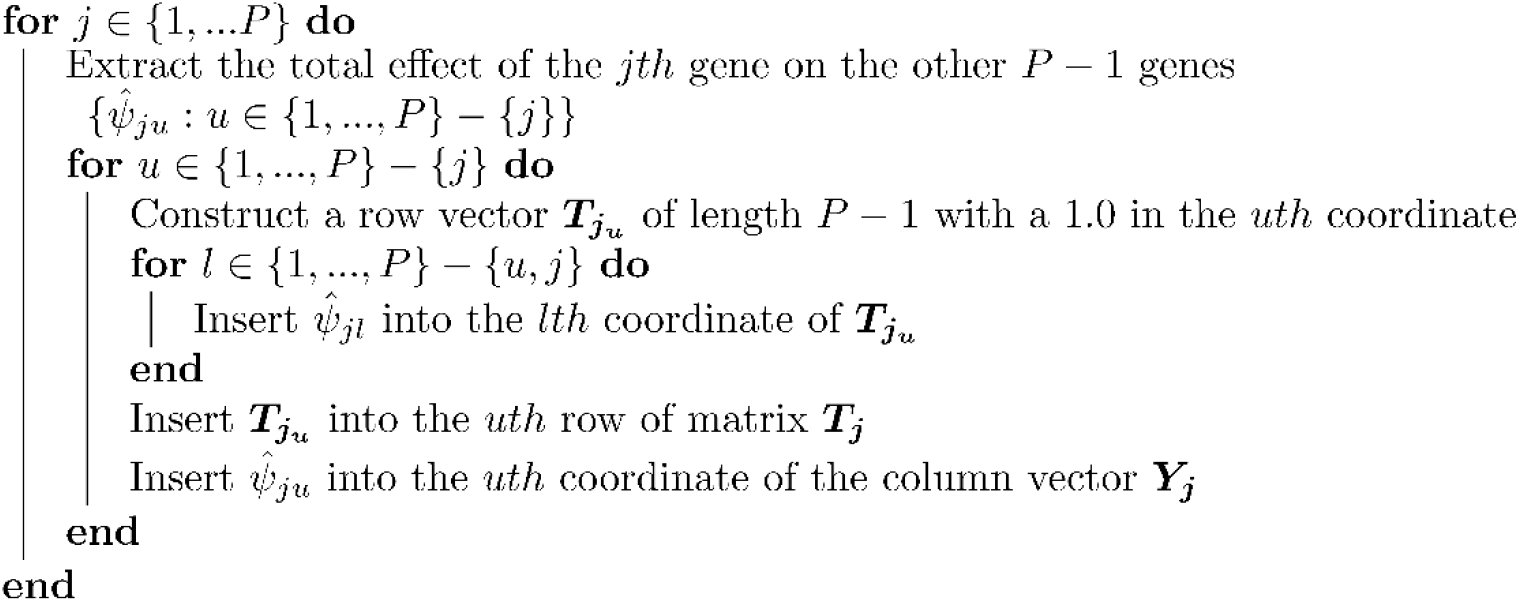

This procedure results in *P* matrices ***T***_***j***_ of size (*P −* 1) *x* (*P −* 1) and *P* column vectors ***Y***_***j***_. We then concatenate 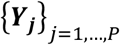 vertically into a column vector ***Y*** of length *P x* (*P −* 1) and we form the block 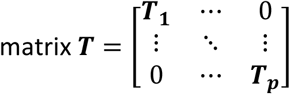

We then define the likelihood as the probability of ***Y*** conditional on ***T*** and parameters ***β*** and σ_*p*_. ***T*** and ***Y*** represent a system of linear equations relating the total effects ***ψ*** to the direct effects ***β***. For a given gene *l* we define the set of rows in ***T*** corresponding to experimental observations where we perturb a putative parent of *l* and record the effect on *l* as 𝕃. For each row of ***T*** and ***Y*** where the *lth* gene is the child node, we specific the likelihood as:

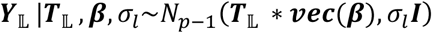

Including a child node specific dispersion parameter *σ*_*l*_ allows for heterogeneity in the residual variance across the genes.

Because we have prior knowledge of what realistic gene networks look like, we specify the prior in three parts as follows:

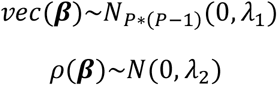

where *ρ*(***β***) is defined as the spectral radius of ***β***, i.e., the maximum eigenvalue of ***β***. We estimate the maximum eigenvalue of ***β*** using power iteration. We incorporate a prior on the spectral radius because it is an upper bound over the NOTEARS DAG penalty^20^, which is a differentiable penalty that enables DAG search in a continuous optimization framework. Importantly, we encode this prior as a “soft-constraint” with the Gaussian density to weakly penalize the divergence of β from the space of DAGs while still allowing for cyclic elements.

Over the columns of ***β***, i.e., ***β***_\****j***_ we place a sparsity inducing L1 prior:

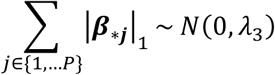

The purpose of this term in the prior is to reflect our belief that the indegree of a gene should be relatively small; we know that genes are not directly regulated by hundreds of TFs. In contrast, a given TF may regulate hundreds of downstream genes, so we do not penalize the rows of ***β***. Overall – this prior encodes the following three prior beliefs: 1. The effects should be somewhat small on a partial correlation scale; 2. The maximum eigenvalue should not be very large to penalize graphs with many cycles; 3. The indegree for each gene should be relatively small, while the outdegree should not be penalized.

On the dispersion terms,σ_*p*_, we place a *LogNormal*(*−*3, 5) prior. We estimate *P* total dispersion terms because there may heterogeneity in the residual variance of the total effects across the KO’d genes.

### Causal network posterior inference

We use pathfinder^30^ to estimate the posterior. Briefly, pathfinder is a variational inference algorithm that optimizes the joint log probability of the model using L-BFGS, i.e., the maximum a posteriori objective. Along this optimization trajectory, it constructs a surrogate posterior at each point using the estimate of the hessian from L-BFGS as the precision of the surrogate posterior. Then, at each point, the evidence lower bound (ELBO) is evaluated. The variational approximation resulting in the largest ELBO is then returned as the posterior estimate. We compute seven runs of this optimization procedure in parallel, and then use importance resampling to combine the fits. We initialize β based on the component-wise sum of the MLE estimate of ***β*** and a vector of gaussian noise i.e. ***β*_*init*_** = 0.1 * ***β*_*MLE*_** + 0.1 * ***z, z*** ∼ *N*(0, 1).

### Causal network posterior uncertainty quantification

We compute a pseudo-posterior inclusion probability (PIP) we defined as *PIP*(*β*_*ij*_) = *P*(|*β*_*ij*_| > *ϵ*). We set *ϵ* = 0.05. We also computed local-false sign rates (LFSR) estimates: *LFSR*(*β*_*ij*_) = min(*P*(*β*_*ij*_ > 0), *P*(β_*ij*_) < 0). We note that these summary statistics, although likely proportional to the ‘true’ values, are likely somewhat uncalibrated given that a) we do not model the underling discrete graph structure *G* separately from the parameters *β* and b) calibrated inference in a network setting has been shown empirically to be extremely challenging.

### Simulation of a cyclic network in a steady state

We start by simulating a given expression vector of *P* genes as ***X*_0_** ∼ *LogNormal*(1.00, 0.10). Then, for a given adjacency matrix ***β*** we model the effect of a perturbation on the *kth* gene as setting ***β***_⋆*k*_ = 0, i.e., we remove the incoming edges to this node. We denote this perturbed adjacency matrix as 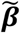 We then sample the “steady-state” limit as 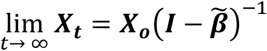 We assessed the performance of our algorithm on a ***β*** corresponding to a cyclic network.

### ABC-GRN

We extracted the CD4+ enhancer to gene predictions from the ABC model^34^ and we intersected them with the differential ATAC peaks from Freimer et al., which were generated on samples where the 24 *IL2RA* regulators were KO’d. For the *ith* gene we included *i* → *j* as an edge in this graph if one its differential ATAC peaks intersected with an ABC enhancer for gene *j*, suggesting that perturbation of gene *i* was perturbing a cis regulatory element for gene *j*. We then calculated the enrichment of these edges among those detected in the *IL2RA* regulator sub-network of causal network estimate.

### HBase validation network

We downloaded the HumanBase^35^ predicted “T-Lymphocyte” network from https://hb.flatironinstitute.org/download. We downloaded the version of the network with only the top edges included. We then estimated enrichment in the same manner as with the ABC-GRN network.

### Bipartite graph model of downstream gene expression

We refer to a “downstream” gene as those that were measured among the 12,803 genes that were highly expressed but not among the perturbed genes. We form a matrix ***Y*** with 12,803 columns containing the vst normalized gene expression data. We define a matrix ***X*** with the expression values of the 84 perturbed genes. We applied the same normalization data procedure as in our causal network estimation such that both ***X*** and ***Y*** are vst transformed data that is orthogonal to covariates (donor identity, PCs 1-4). We specified the following likelihood for the *ith* measurement of the *jth* downstream gene:

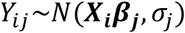

Over the ***β***_***j***_ we place the following prior *β*_***j***_ ∼ *N*_*p*_(0, *α* * Σ_*β*_), where Σ_*β*_ is defined as the asymptotic steady state covariance implied by our point estimate from the causal network model, i.e., 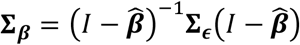. This prior encodes the belief that similar effects among the 84 genes in the causal network will increase the likelihood of similar downstream effects. Because we used a conjugate prior the posterior has an analytic form:

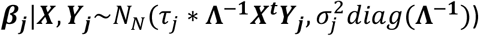

where Λ = (Σ_*β*_ + *τ*_*j*_ * ***X*^t^*X***) and 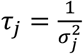

We set α = 0.10 in practice, although in principle empirical Bayesian approaches or other criteria could be used to set this hyperparameter. We estimate the residual variance parameter 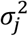 using maximum likelihood and we use lfsr as our variable selection criteria.

### Pathway analysis

Downstream enriched pathways were identified for each perturbation using pathfindR (v1.6.4)^67^. For each upstream gene perturbed, outgoing edges within the BG model were used as input for pathfinder, with a significance threshold of LFSR < 5 x 10^−3^. Gene sets were limited to KEGG^40^, Reactome^45^, and GO-BP^50^ and the minimum gene set size and enrichment threshold were set to 10 and 0.05 respectively. Pathways were prioritized for visualization based on the number of genes within the module with enrichment for the pathway, median fold enrichment across all members of the module, and relevance to T cell biology.

### LD score regression analyses

We first defined gene sets corresponding to each of the nine modules (1A-C, 2A-C, 3A-B, 4) and module 0, which we defined as the set of genes that were expressed highly enough for analysis but were not associated with any of the KO’d genes (at a LFSR threshold of 5 x 10^−3^). For each of these 10 gene sets, we then linked SNPs to these genes (S2G) using seven possible methods following Dey at al^68^, including approaches that link SNPs based purely on physical distance to the nearest gene, fine-mapped eQTLs, promoter Hi-C capture, the ABC model, among others.

For each of the 10 phenotypes analyzed (Supplementary Table S7) we obtained the GWAS summary statistics and performed LD score regression analysis. We included the LD score baseline model v2.1 in the regression. We used the publicly available European ancestry LD score estimates for the HapMap SNPs available from: gs://broad-alkesgroup-public/LDSCORE/Dey_Enhancer_MasterReg/processed_data.

### ATAC and ChIPseq data visualization

Bigwigs for each of the tracks were downloaded from ChIP-Atlas. ATAC bigwigs and differentially expressed regions were procured from Freimer et al. and a representative donor was used for visualization of each perturbation effect at the IL17F locus. Visualization was performed with rtracklayer (v1.52.1) and ggplot2 (v3.4.1). APRIS gene structure was used for gene annotation with gggenes (v0.5.0).

We included data from the following SRA sources:

STAT5B KO ATAC-SRX10558086, KMT2A KO ATAC-SRX10558079, AAVS1 KO ATAC-SRX10558063, H3K4me3 ChIP-activated Th17 ChIP-SRX16500373 (GSM6376841), STAT5B ChIP-SRX041293 (GSM671402)

## Extended Data Figures

**Extended Data Figure 1.**
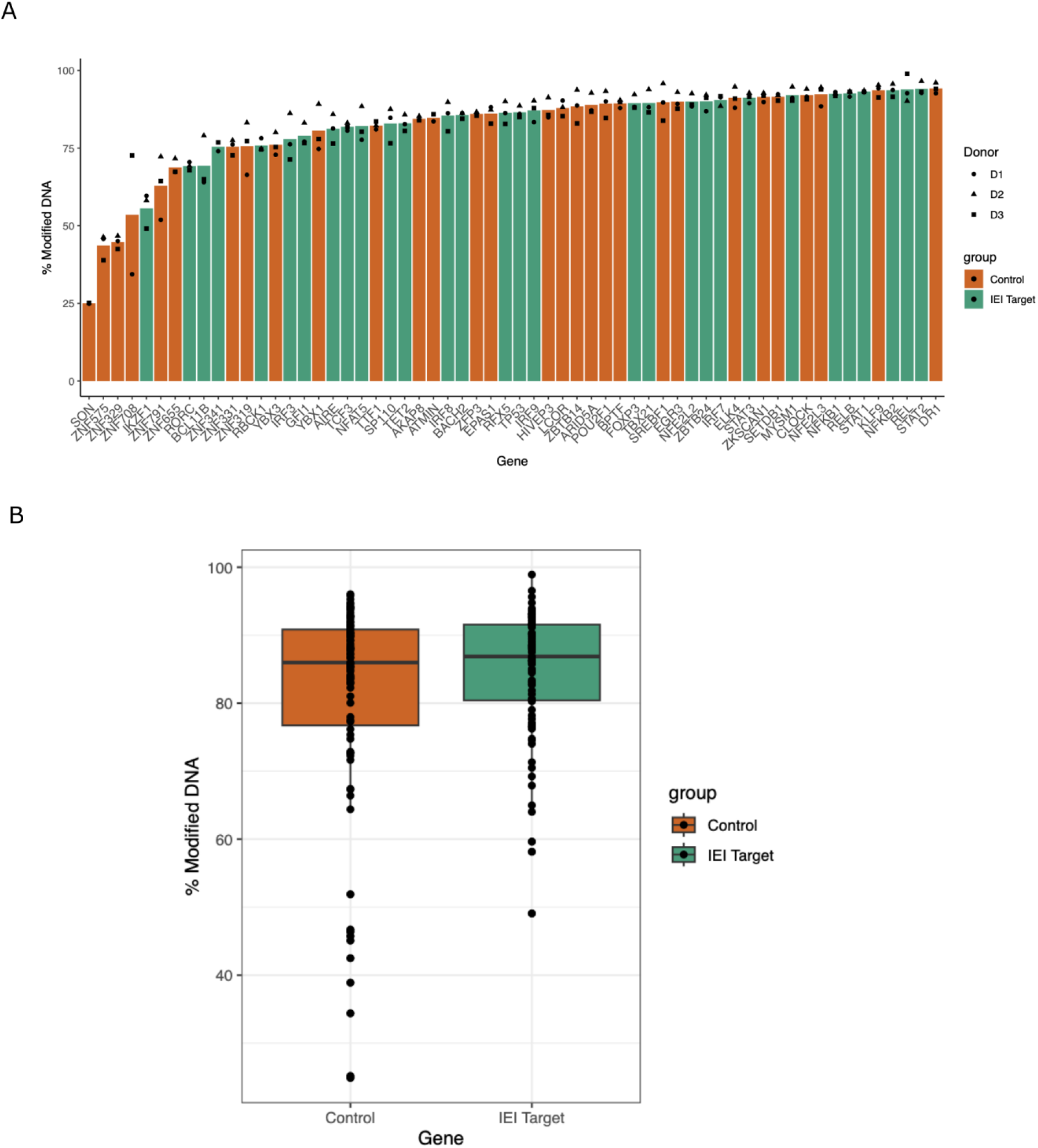
CRISPR editing efficiency by gene group. **A** Percent of reads with indels, stratified by individual gene. **B** Percent of reads with indels, aggregating by gene group.

**Extended Data Figure 2.**
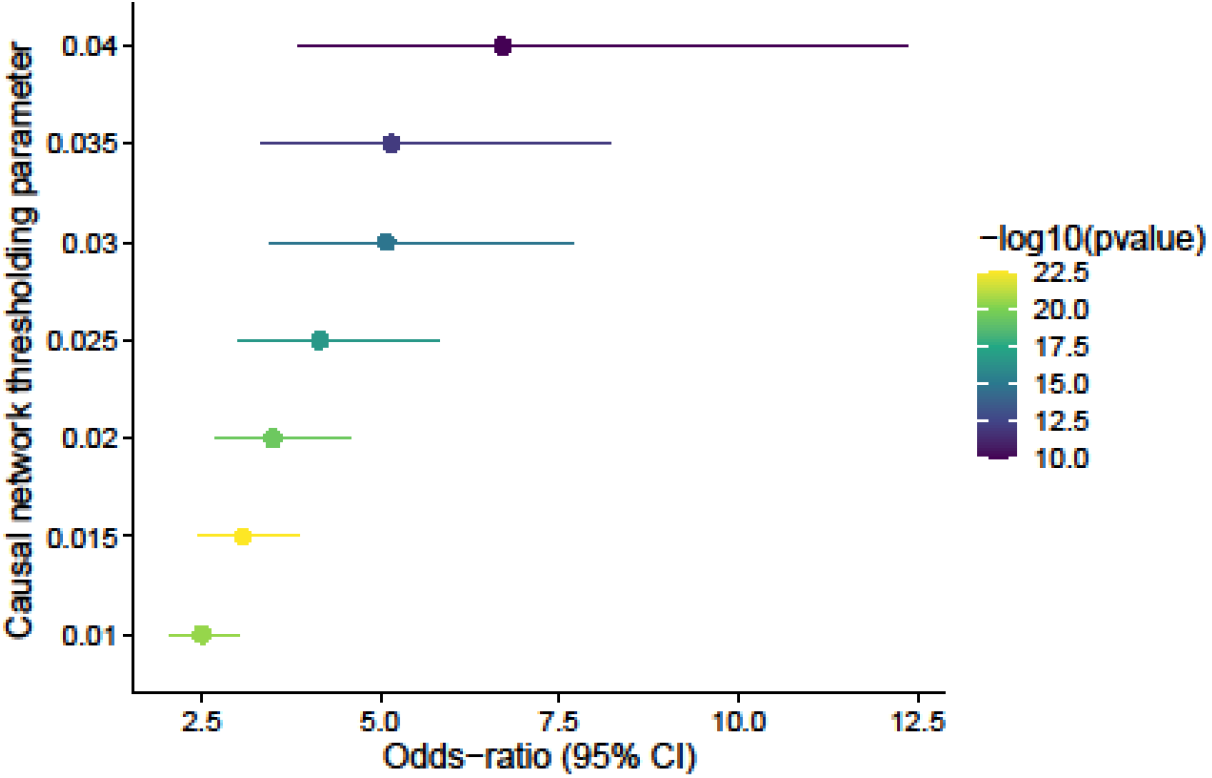
Enrichment of LLCB posterior mean edges in the ABC-GRN validation network.

**Extended Data Figure 3.**
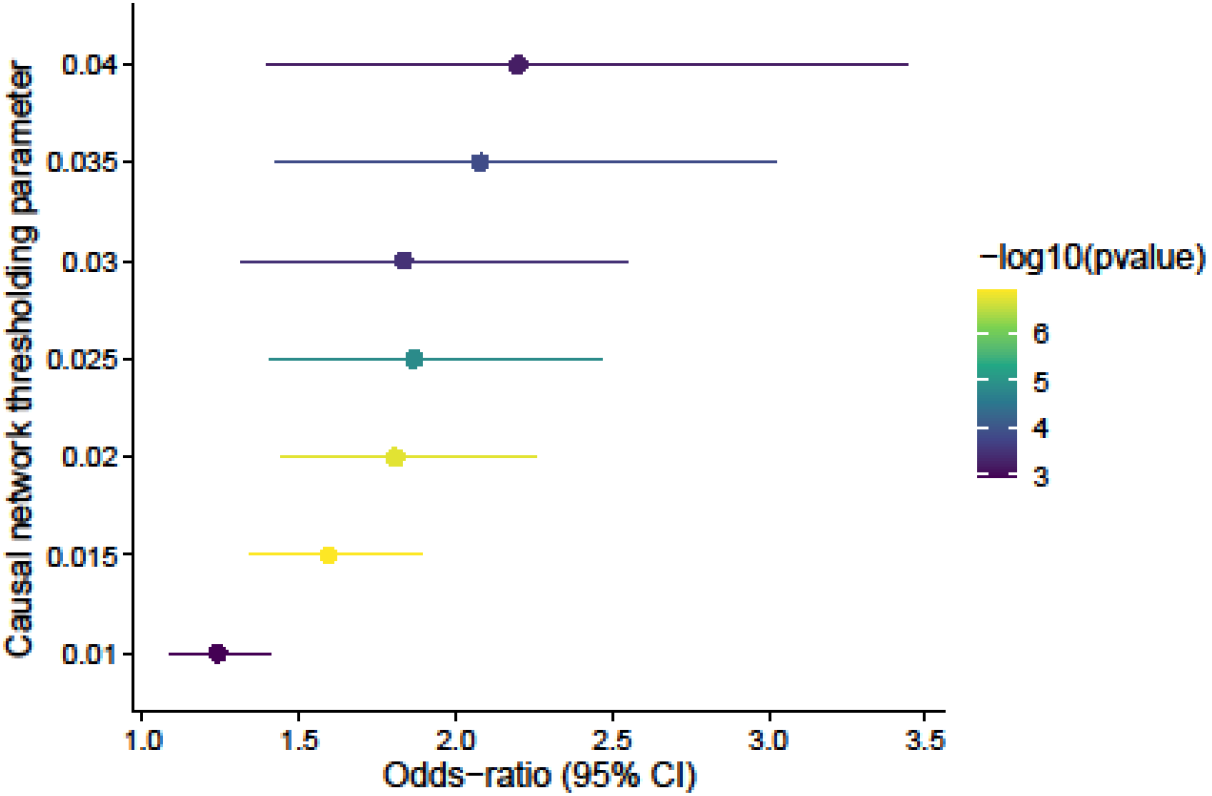
Enrichment of LLCB posterior mean edges in the HBase T-cell network.

**Extended Data Figure 4.**
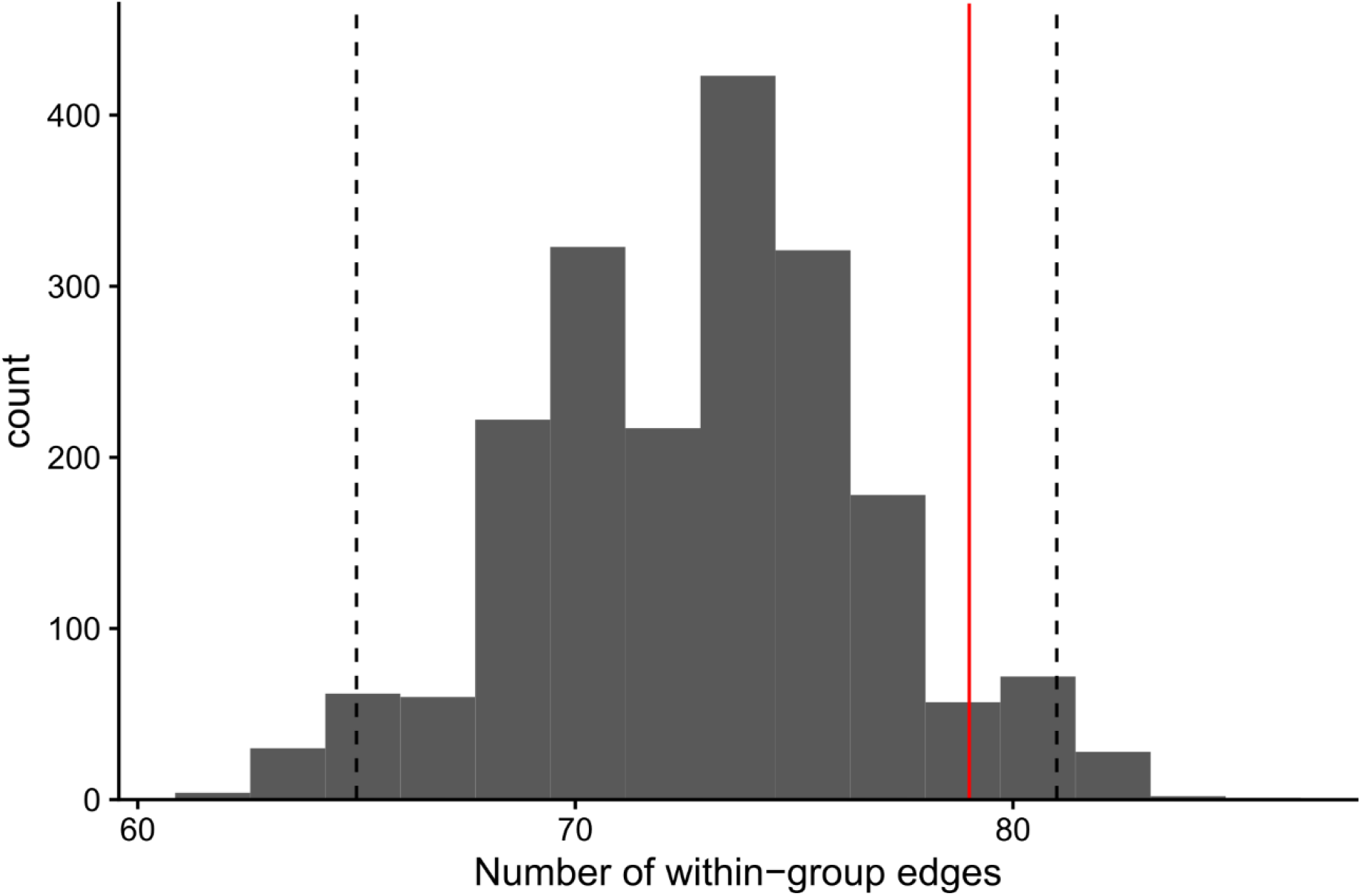
Permutation test of the number of edges between genes that share the same gene group. A set of 2,000 null permutations of the network were generated by using the rewiring algorithm to preserve the node degree. Within each permutation, the number of edges with the same gene group were counted. The observed value is denoted by the red vertical line, and the empirical 2.5% and 97.5% quantiles from the permuted data are denoted by vertical dashed lines.

**Extended Data Figure 5.**
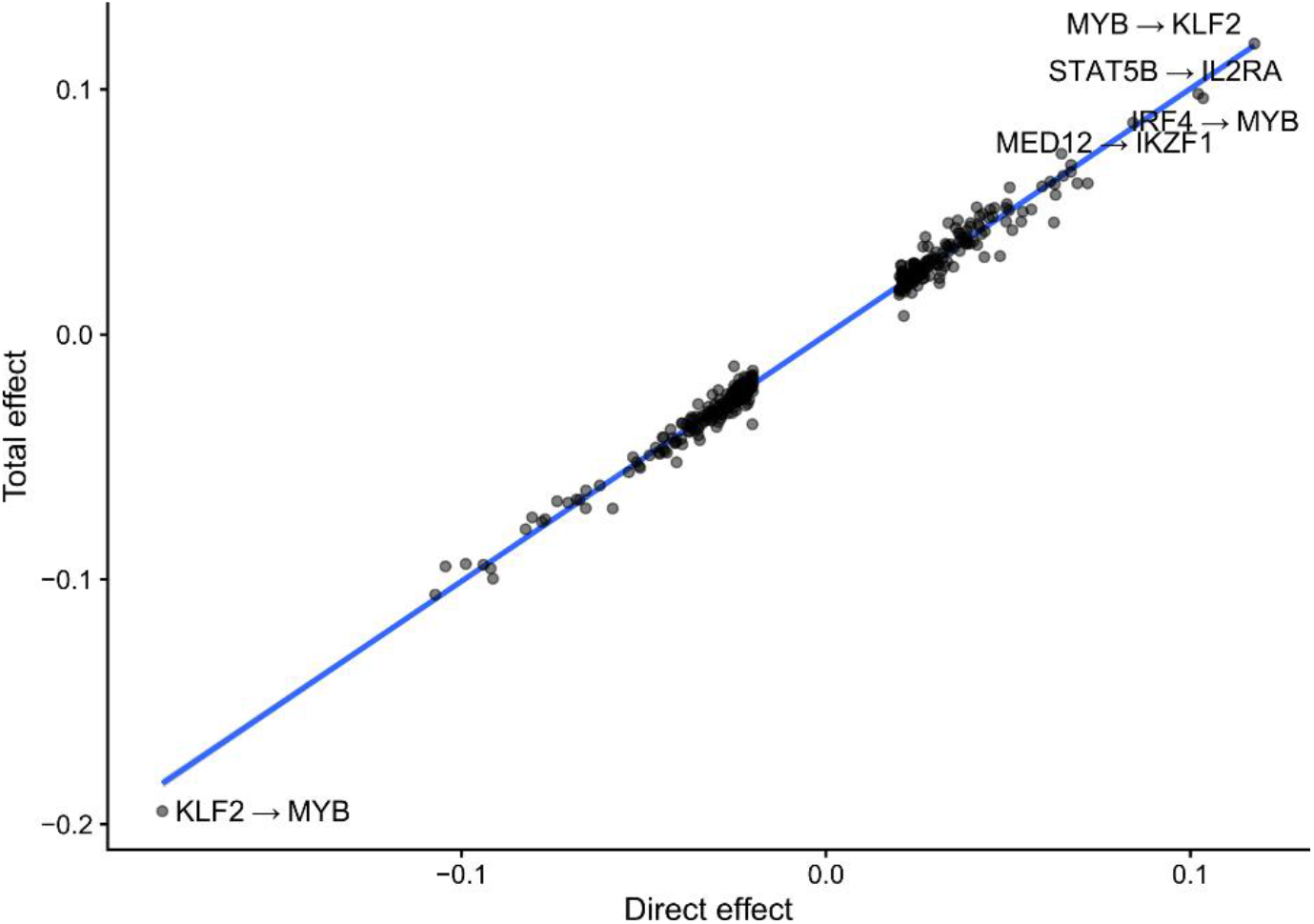
Comparison of direct to total effects among the 84 KO’d genes. The x-axis is defined as the posterior mean estimates of the adjacency matrix estimated by LLCB. Units are in terms of standard deviations of normalized gene expression. The y-axis is estimated through the processing procedure described in Methods.

**Extended Data Figure 6.**
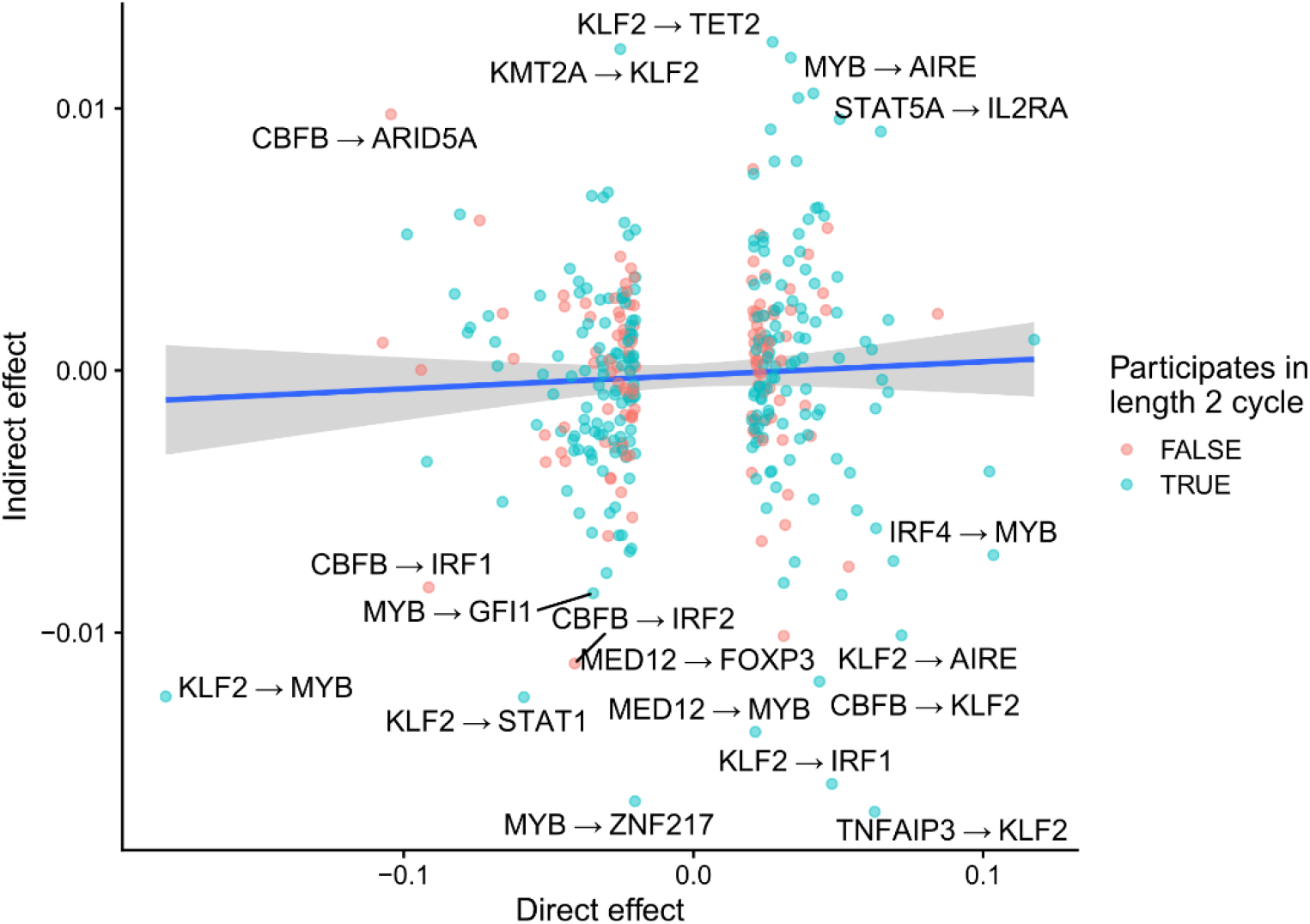
The largest indirect effects are mediated by cycles of short length.

**Extended Data Figure 7.**
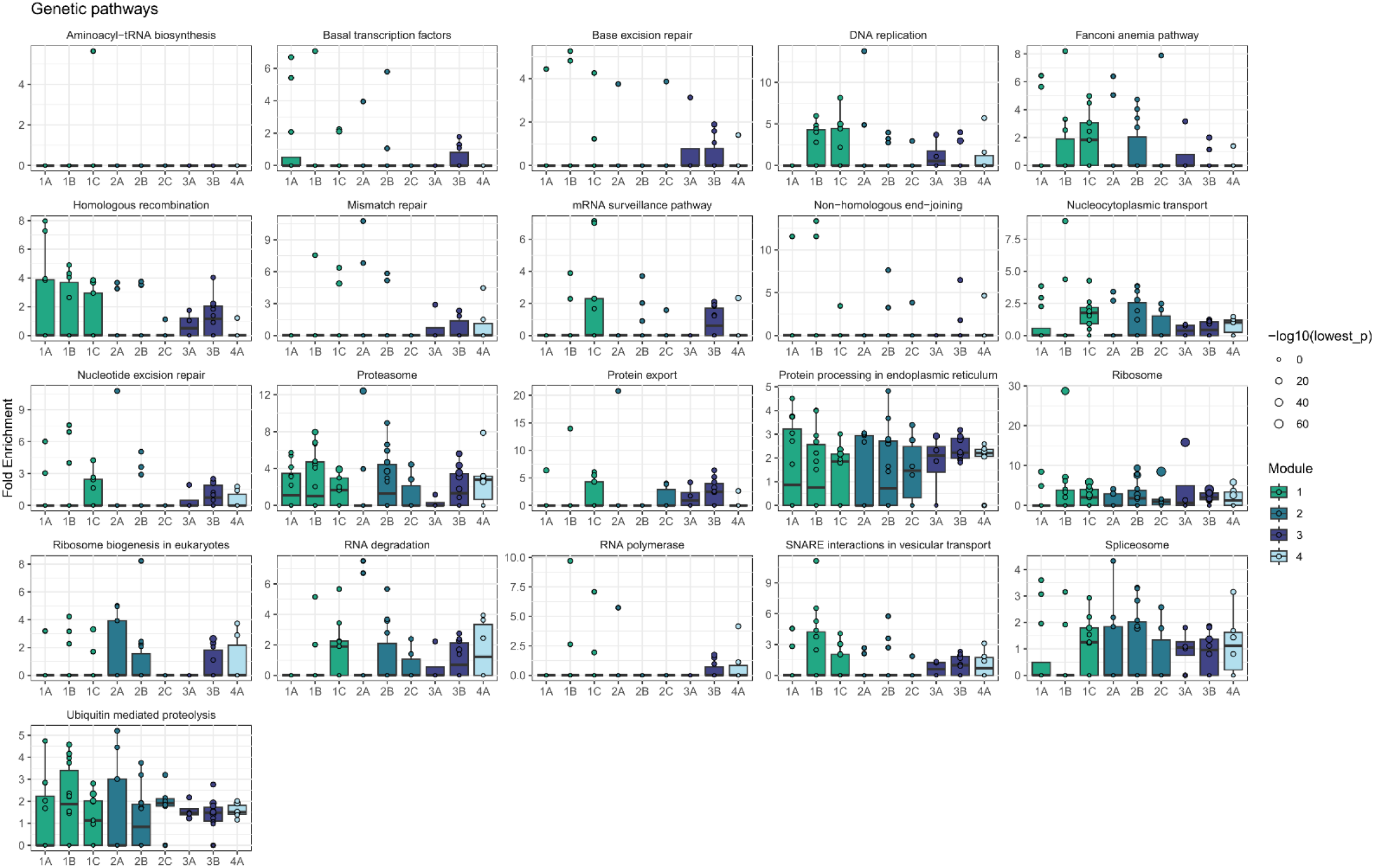
Enrichment of module effects on KEGG signaling pathways. Enrichment analyses were performed with pathfindR.

**Extended Data Figure 8.**
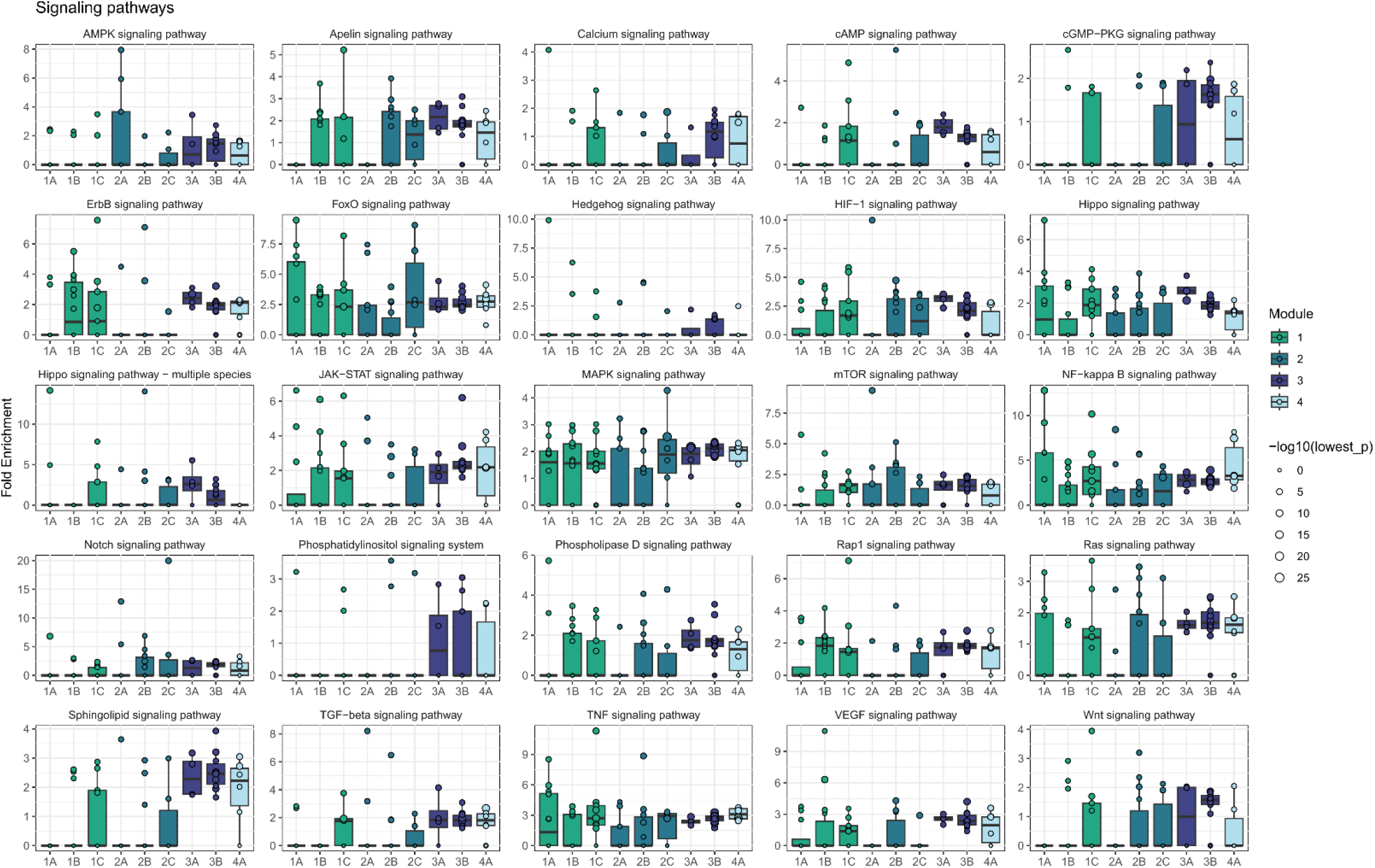
Enrichment of module effects on KEGG signaling pathways. Enrichment analyses were performed with pathfindR.

**Extended Data Figure 9.**
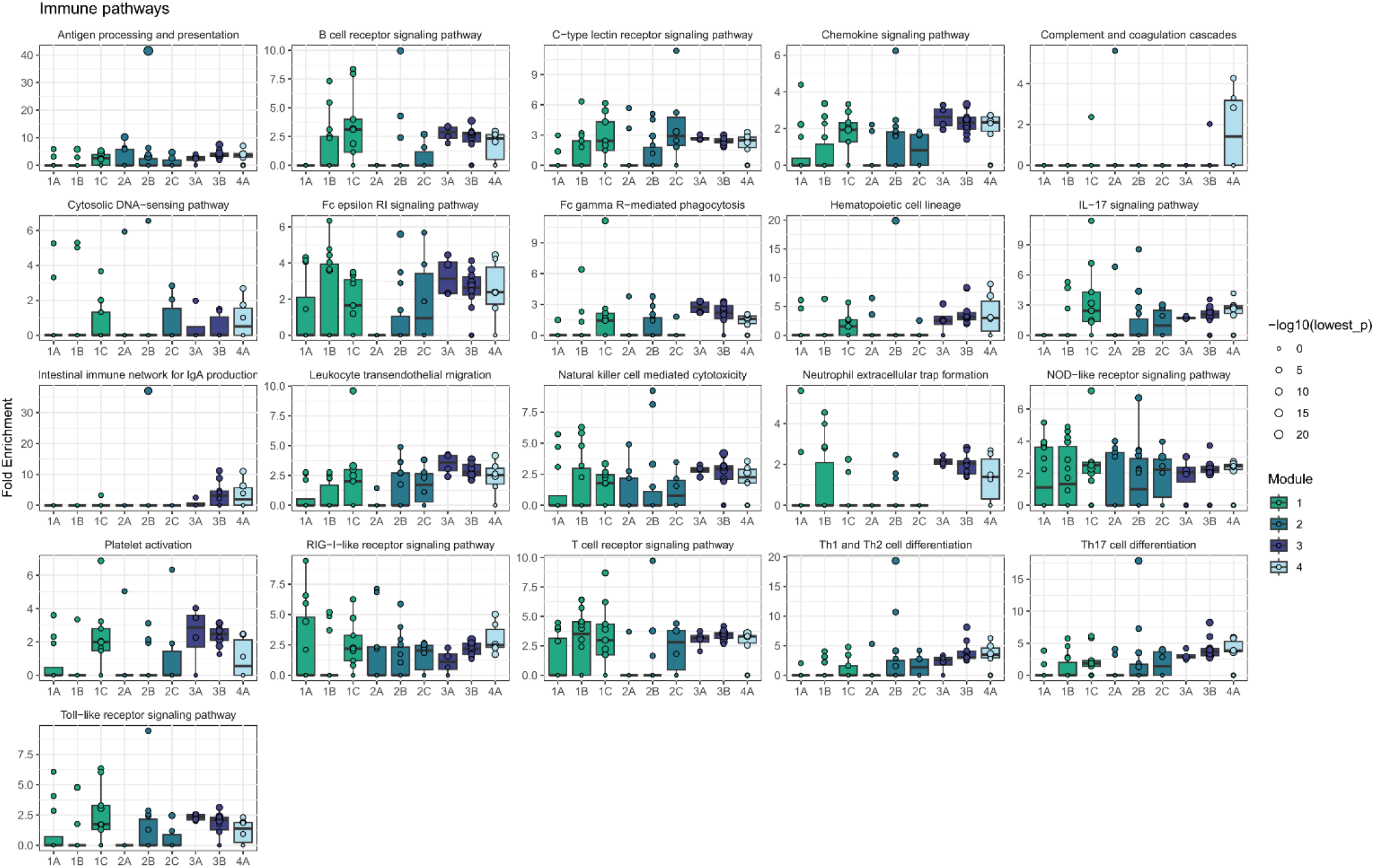
Enrichment of module effects on KEGG immune pathways. Enrichment analyses were performed with pathfindR.

**Extended Data Figure 10.**
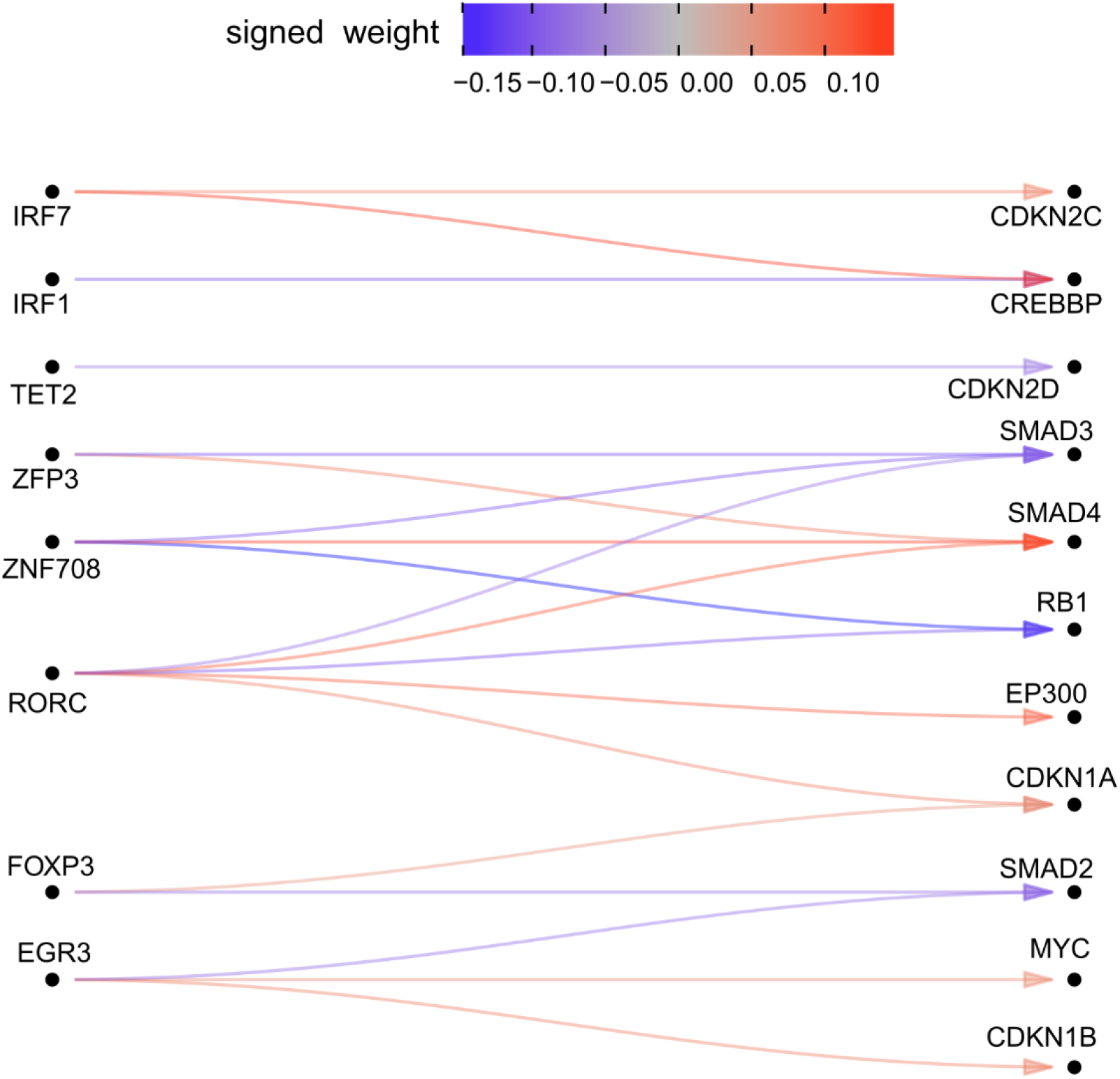
Network plot demonstrates the effect of the cluster 2A upstream regulators on cell-cycle genes. The network using edges estimated from the BG model are plotted. Colors indicate the effect size and arrows indicate the direction of effect. The genes on the left-hand side are among the 84 KO’d genes, and the genes on the right are genes that are listed among the KEGG cell cycle pathway genes.

**Extended Data Figure 11.**
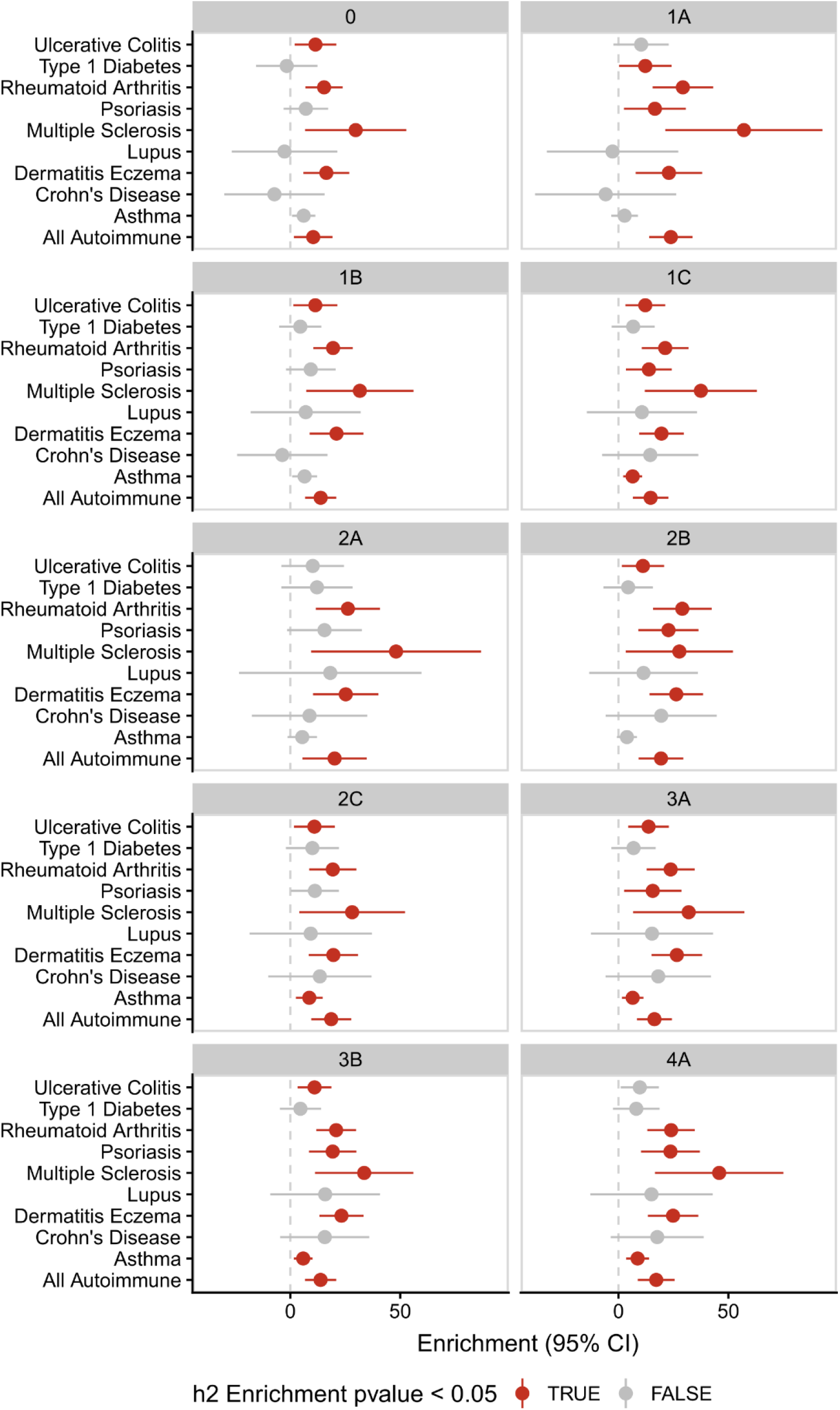
Marginal heritability estimates from LD score regression. LD score regression was used to estimate the heritability enrichment of SNPs linked to genes in each module for each phenotype. SNPs were linked to genes using the ABC predictions in T cells.

**Extended Data Figure 12.**
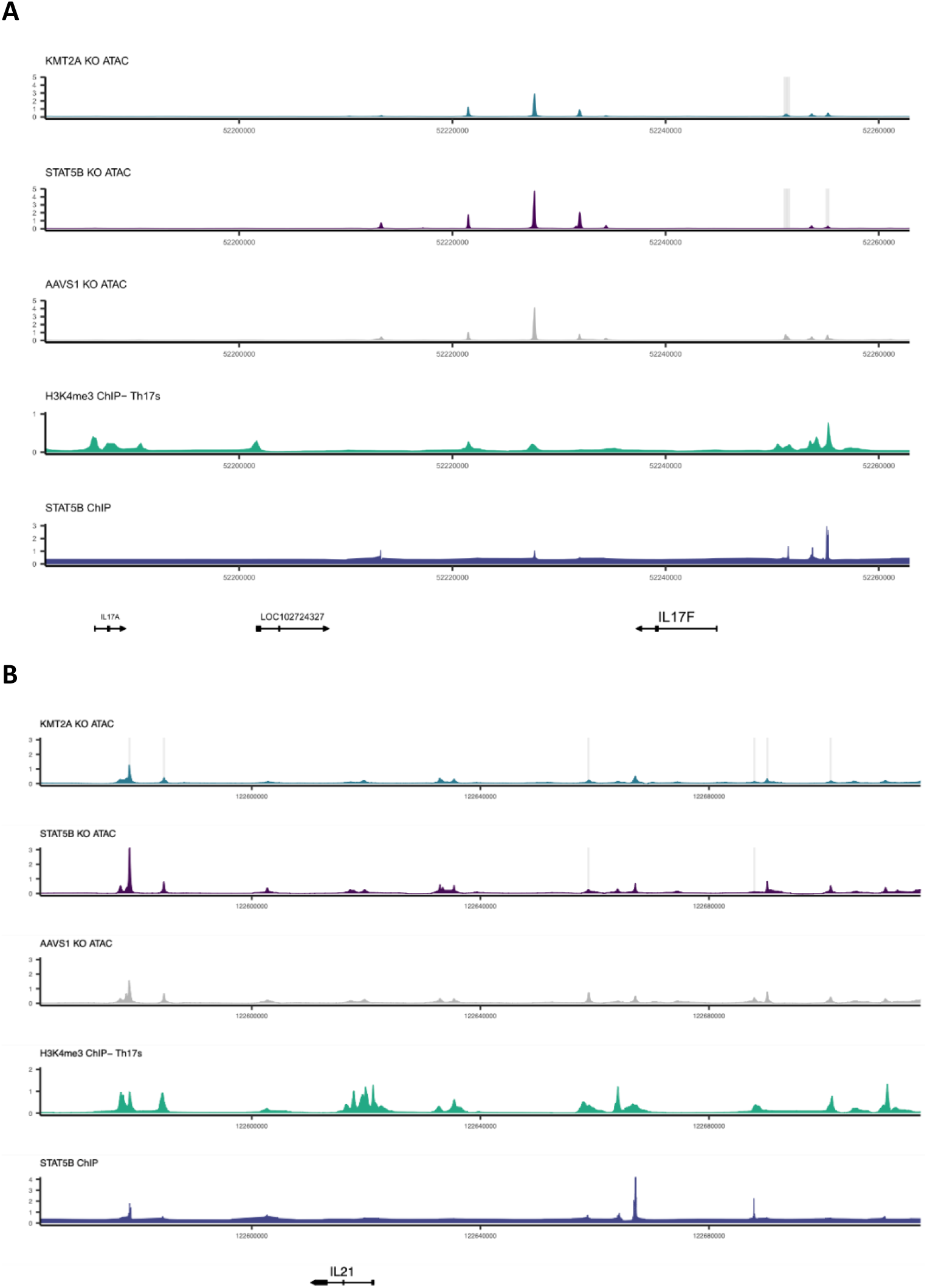
KMT2A and STAT5B jointly regulate chromatin accessibility at the *IL17F* locus (A) and *IL21* locus (B).

**Extended Data Figure 13.**
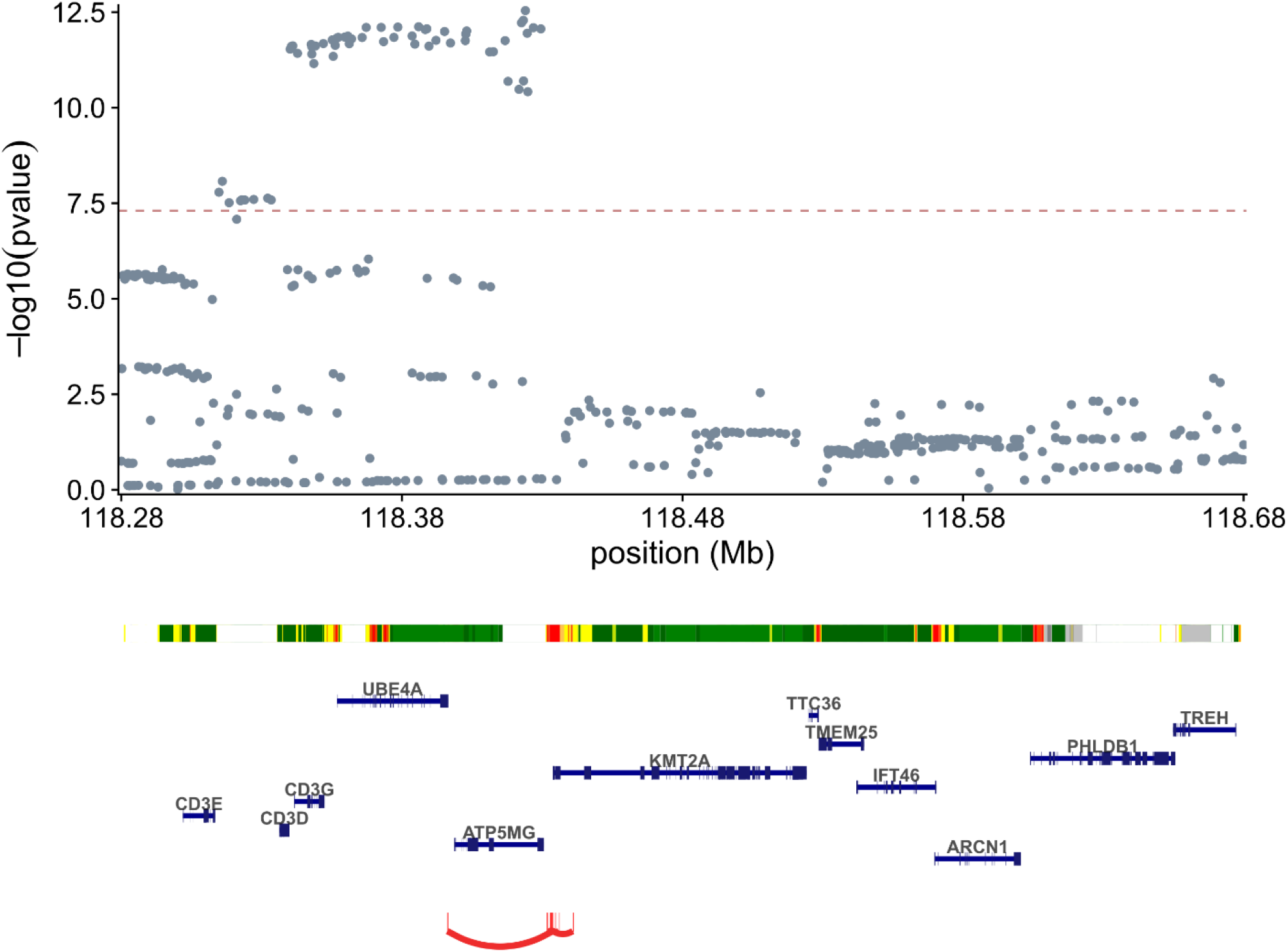
Meta-analysis of autoimmune GWAS from Shirai et al. and Finngen v8. The *KMT2A* locus plot is displayed with a chromHMM^69^ track from Th17 cells. The predicted enhancers of *KMT2A* from the ABC model in CD4+ T cells are shown in red arcs at the bottom.

## Acknowledgements

We thank Romain Lopez, Matt Aguirre, Eric Kernfeld, and Prashanthi Ravichandran for several stimulating conversations about gene networks and structure learning. We also thank members of the Pritchard, Battle, and Marson labs for feedback and insightful discussions. M.M.A. is a National Science Foundation Graduate Research Fellow supported under Grant No. 2038436. J.W.F. was funded by NIH grant R01HG008140. M.O. was supported by Astellas Foundation for Research on Metabolic Disorder and Chugai Foundation for Innovative Drug Discovery Science (C-FINDs). J.P., M.O., and A.M. are supported by the NHGRI (2R01HG008140). A.M. received funding from the Simons Foundation, Career Award for Medical Scientists, a Lloyd J. Old STAR Award (Cancer Research Institute), Parker Institute for Cancer Immunotherapy, Innovative Genomics Institute. A.B. was supported by NIGMS R35GM139580.

## Code Availability

https://github.com/weinstockj/RNAseq-perturbation-CD4-pipeline

https://github.com/weinstockj/LLCB

## Data Availability

Raw sequencing from Freimer et al. was previously deposited at GEO under accession GSE171737.

